# A secreted endosymbiont protein essential for colonizing host cells

**DOI:** 10.1101/2025.12.01.691566

**Authors:** Gerald P. Maeda, Allen Z. Xue, Ethan W. Yu, Aadhunik Sundar, J. Elijah Powell, Thomas E. Smith, Nancy A. Moran

## Abstract

Intracellular bacterial symbioses have arisen myriad times in eukaryotes, with dozens known from insects alone^1,2^. Beginning with *Buchnera*, the obligate endosymbiont of aphids, genomes of endosymbionts have illuminated their evolutionary origins and metabolic contributions to hosts^3,4^. However, the mechanisms by which nonculturable endosymbionts enter host cells and suppress cellular immune processes have remained unknown. We show that an uncharacterized *Buchnera* protein, here designated SyeA, was present in the *Buchnera* ancestor, is secreted and homologous to secreted effectors of bacterial pathogens and is essential for *Buchnera* transmission. *Buchnera* is transmitted via expulsion from specialized maternal cells and uptake by embryos^5^. Using immunofluorescence microscopy, we found that SyeA levels peak upon colonization, accompanied by actin accumulation at the entry site. SyeA localizes outside the host-derived membrane and actin layer surrounding each *Buchnera* cell and colocalizes in host cytoplasm with Rho1, which regulates actin polymerization. *syeA* knockdown disrupts colonization and embryonic development and elevates lysosomal activity, leading to *Buchnera* destruction^6^. Our findings provide rare insight into how an anciently associated, mutualistic endosymbiont achieves its intracellular existence. SyeA is a vestige of pathogenic origins followed by evolution of increased host control and erosion of the original, more complex pathogenicity machinery.

## MAIN TEXT

As for many obligate endosymbionts, *Buchnera* benefits hosts by provisioning essential nutrients, and aphids require *Buchnera* for successful embryonic development^7^. Despite its reduced genome, *Buchnera* retains pathways for producing amino acids needed by hosts^3^. *Buchnera* cells densely populate the cytoplasm of bacteriocytes, which are specialized host cells formed during embryogenesis^8,9^. Each *Buchnera* cell is housed in a cytoplasmic compartment, the symbiosome, enclosed within a host-derived membrane. Transmission occurs when *Buchnera* cells are exocytosed from maternal bacteriocytes and endocytosed by a large multi-nucleate embryonic cell, which develops into mature bacteriocytes^5^. In older aphids, symbiosomes fuse with lysosomes, resulting in death of the resident *Buchnera*, hypervacuolation of the cytoplasm, death of the bacteriocyte itself, and recycling of the released nutrients^6,10^.

To find clues as to how *Buchnera* achieves its intracellular existence, we sought to identify endosymbiont molecules that interface directly with hosts, such as outer membrane proteins or secreted proteins. Despite genome reduction, *Buchnera* of the pea aphid *Acyrthosiphon pisum* (hereafter *Buchnera*-Ap) retains an apparent secretion system, as it possesses 26 genes encoding components of the flagellar basal body though not the flagellar filament itself^3^. The basal body is homologous to, and can function as, a Type 3 Secretion System (T3SS) that delivers proteins across the bacterial cell wall and into host cells^11^. In *Buchnera-*Ap, the basal body components are highly expressed^12–14^ and number in the hundreds on the *Buchnera* cell surface^15^ (Extended Data Fig 1).

Secreted effectors are candidates for *Buchnera* molecules that interact with host cells and enable the intracellular lifestyle, but their identities have remained unknown. We searched *Buchnera* proteomes for candidate effectors, leading us to focus on an uncharacterized gene encoding a hypothetical protein with few homologs outside *Buchnera.* We thus undertook efforts to better characterize this protein and its role(s) in the symbiosis.

## RESULTS

### A secreted protein in diverse *Buchnera*

To identify proteins secreted by *Buchnera*’s basal bodies, we analyzed sequences for all 573 chromosomally encoded proteins of *Buchnera*-Ap (str. APS, GenBank accession NC_002528) using BastionX, which scans for signatures of T3SS effectors^16^ (Dataset S1). Eight proteins were predicted as potential effectors (Table S1). The top-scoring candidate was an uncharacterized protein previously referred to as Yba3 (UniProt P57644), and here designated SyeA, for “SYmbiosis Essential”, based on evidence presented below. Other predicted effectors included SyeB (previously Yba4), a short uncharacterized protein cotranscribed with SyeA (Fig 1b)^17^, and the flagellar hook protein (FlgK) which is secreted during basal body assembly.

**Fig 1.**
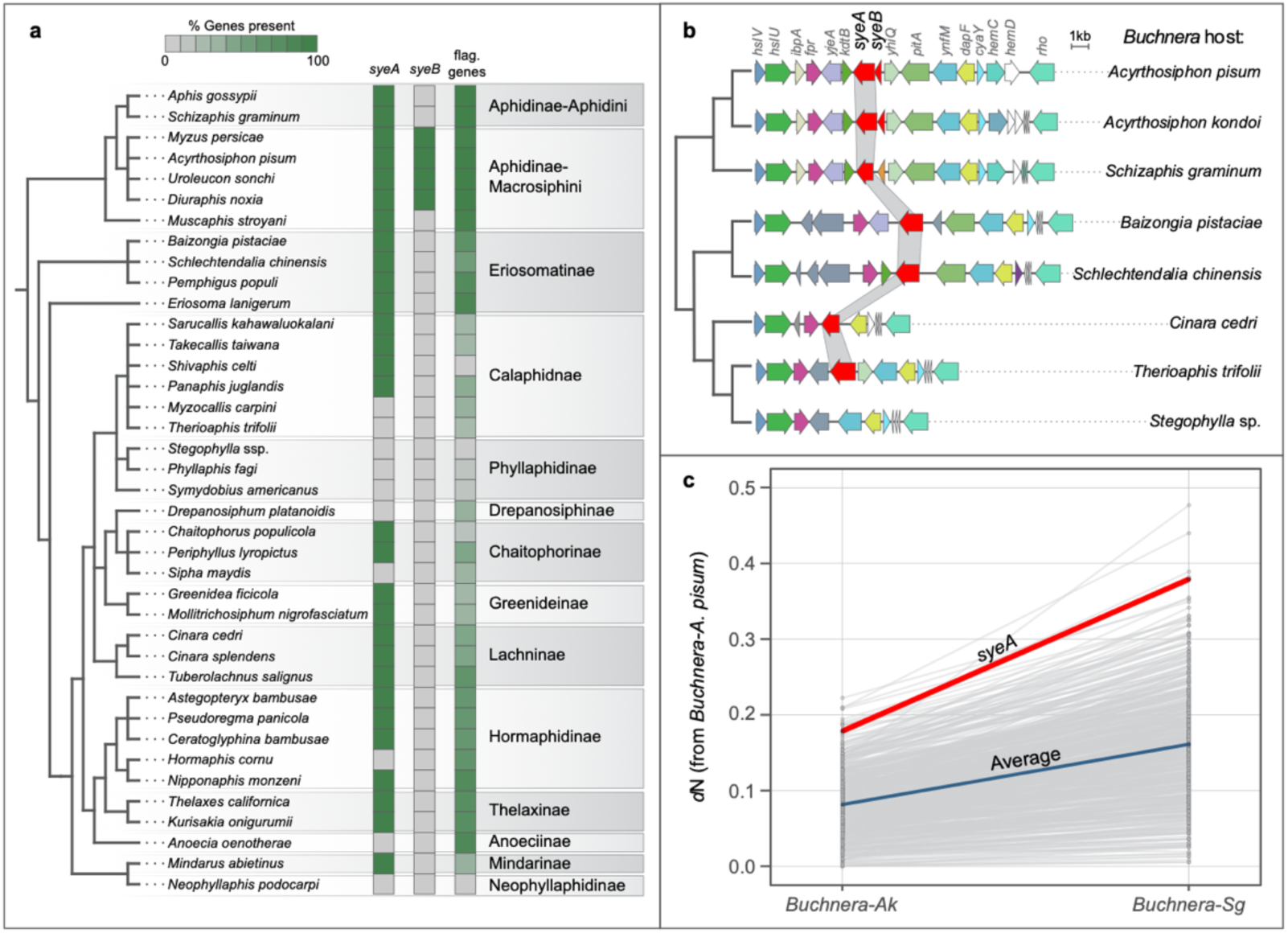
Evolution of *syeAB* in *Buchnera.* **a**) Presence of *syeA*, *syeB*, and genes for flagellar basal body components across *Buchnera* from diverse aphid hosts. The *Buchnera* phylogeny corresponds to a published analysis^50^ with less-supported nodes collapsed. Aphid subfamilies for each host species are included on the right, with tribes included for the Aphidinae. **b**) Retention, loss, and gene neighborhood of *syeA* across representative *Buchnera* lineages. **c**) Nonsynonymous divergence of shared *Buchnera* proteins for pairwise comparisons of *Buchnera*-Ap to *Buchnera*-*Acyrthosiphon kondoi* and *Buchnera*-Ap to *Buchnera*-*Schizaphis graminum*. *d*N was calculated for 526 protein-coding genes shared across the three *Buchnera* genomes. The average (arithmetic mean) for all shared proteins is shown in blue, *syeA* in red, and all other proteins in gray.

In the initial report for the *Buchnera-*Ap genome*, syeA* and *syeB* (GenBank protein IDs BAB13273, BAB13274) were among 4 orphan genes (i.e., having no detectable homologs in current databases)^3^. Of the other two, one (*yba1*) reflected an annotation error, and the other (*yba2*, GenBank ID BAB12898) was subsequently found to have homologs in numerous bacteria^17^.

We surveyed 115 available *Buchnera* genomes for the presence of *syeAB* and genes for basal body components, using OrthoFinder^18^ (Datasets S2, S3). Because *syeA* and *syeB* evolve rapidly, we also manually inspected the relevant genomic neighborhood to detect divergent homologs (Fig 1b).

*syeA* homologs were detected in 103 of the 115 *Buchnera* genomes (Fig 1a, Dataset S4). Often, homology was evident only in the C-terminal portion of the protein, corresponding to amino acids 163–367 of *Buchnera*-Ap. The N-terminal portion of SyeA and all of SyeB contain alpha helices and disordered regions (Extended Data Fig 2). Even for the more conserved C-terminal region, amino acid identity was low (∼25%) for distantly related *Buchnera*. *syeB* was recognizable only in *Buchnera* of aphids in the tribe Macrosiphini, which includes *A. pisum*, although a small hypothetical protein encoded upstream of *syeA* is also present in *Buchnera* of Aphidini.

To determine whether SyeA orthologs in diverse *Buchnera* are likely to be secreted, we analyzed them using BastionX and found that all 103 were predicted as T3SS effectors (Table S1, full results in Dataset S5).

Most *Buchnera* retain all or most of the 26 genes for making basal bodies, but some have lost the entire set (Fig 1a, Dataset S6). In the largest aphid subfamily, the Aphidinae, basal body genes and *syeA* are consistently present. In some other aphid lineages, such as Phyllaphidinae, *Buchnera* genomes have lost *syeA* along with some or all genes for basal body components.

To search for homologs outside *Buchnera*, we used SyeA of *Buchnera*-Ap as a query in blastp searches across the GenBank nr protein database, excluding *Buchnera*. SyeA had significant homology in only two groups. Surprisingly, both are intracellular symbionts of aphids or related insects. *Serratia symbiotica* str. Tucson (accession EFW13163,%id = 25.6%, e-val = 6e-06) is an accessory symbiont that sometimes co-resides with *Buchnera*-Ap^19^. *Liberibacter* species (WP182016853, %id = 25.1%, e-val = 5e-05) are endosymbionts of psyllids (sap-feeding insects related to aphids) and plant pathogens^20^. These symbionts are not close relatives of *Buchnera* or each other, implying ancient horizontal transfer of *syeA*.

### SyeA evolves fast but conserves structure

We calculated estimates of pairwise *d*N (nonsynonymous substitutions per nonsynonymous site) for all protein-coding genes between *Buchnera*-Ap, and each of two *Buchnera* strains from other Aphidinae species, *Acyrthosiphon kondoi* and the more distantly related *Schizaphis graminum*. For *syeA*, using values for the alignable C-terminal region only, the rate of protein evolution is among the highest for any gene (Fig 1c). For *Buchnera-A. kondoi and Buchnera-S. graminum* respectively, *d*N values and rankings were 0.179 (12th of 529) and 0.379 (5th of 529).

To examine consequences of amino acid sequence divergence for protein structure, we performed structural alignment of SyeA from divergent *Buchnera* lineages. Structure of the C-terminal 170 amino acids of SyeA from *Buchnera*-Ap aligns with high confidence to the C-terminal region from other *Buchnera*, including highly divergent strains (*Buchnera*-Ua: Zscore = 27.3, RMSD = 0.9, *Buchnera*-Cc: Zscore = 20.6, RMSD = 2.4, *Buchnera*-Bp: Zscore = 21.7, RMSD = 2.0) (Fig 2a, Extended Data Movie 1). Thus, despite extreme sequence divergence, the C-terminal portion of SyeA is conserved at the structural level.

**Fig 2.**
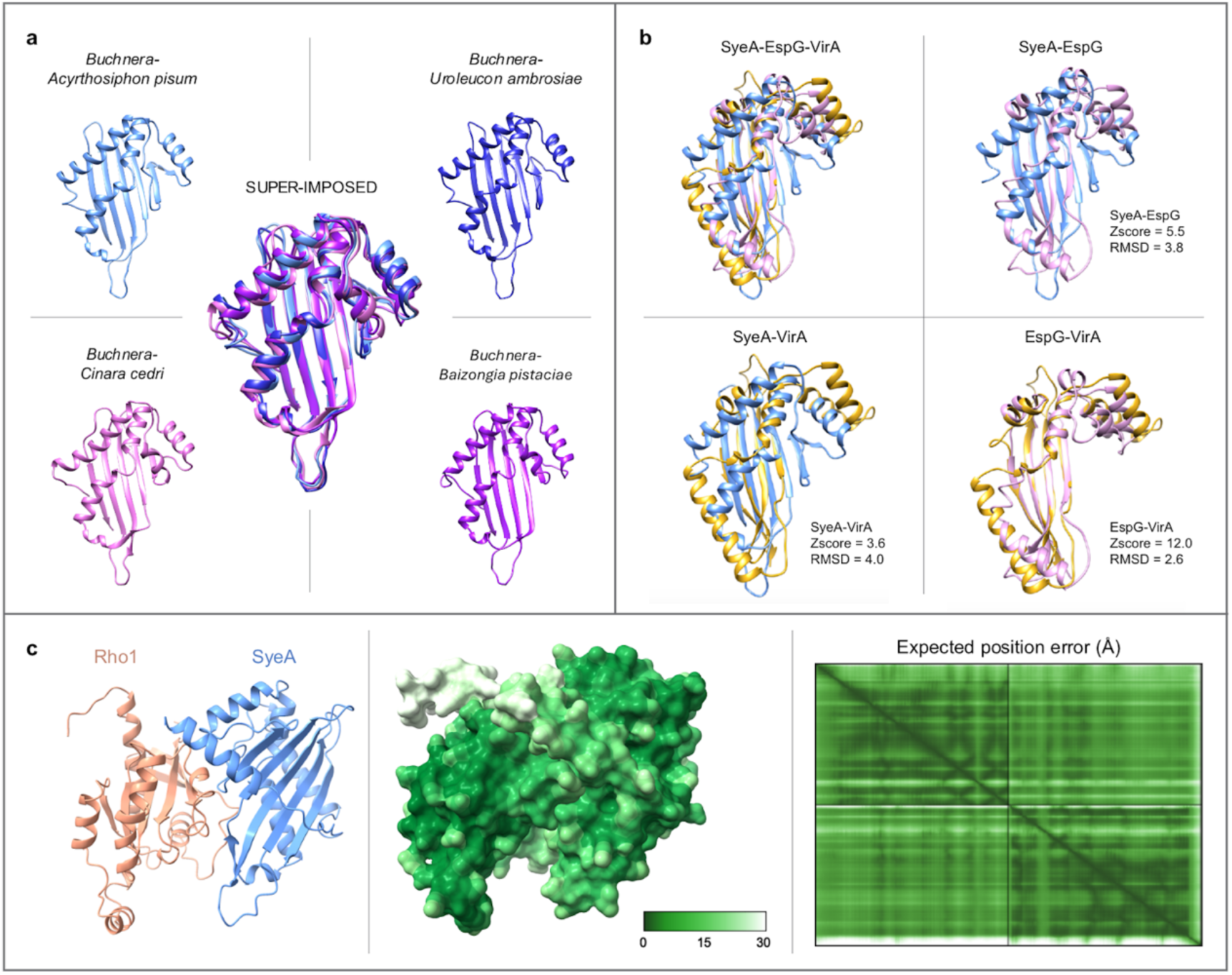
Structural predictions for SyeA homology and interactions. **a**) Structural conservation of C-terminal region of SyeA across divergent *Buchnera* strains. **b**) Structural alignments between C-terminus region of SyeA from *Buchnera*-Ap (blue) with EspG of *Escherichia coli* (pink) and VirA of *Shigella flexneri* (gold). (Z-score and RMSD values are for the C-terminal region only.) **c**) Predicted interactions between C-terminal region of *Buchnera*-Ap SyeA and Rho1 of *A. pisum*. Left: Ribbon depiction of Rho1 (salmon) and SyeA (blue) interaction. Center: Space-filling model of Rho1-SyeA interaction, shaded by prediction confidence as predicted alignment error (PAE) in Ångströms. Right: 2D PAE plot for Rho1-SyeA interaction.

### Structural homology to pathogen effectors

Given this structural conservation, we suspected that structural searches might reveal non-*Buchnera* proteins with similar function. Using the AlphaFold predicted structure for the full-length SyeA of *Buchnera*-Ap, DALI structural searches against empirically determined structures in the Protein Data Bank (PDB), significant structural similarity was evident for EspG of *E. coli* 0157:H7 (Zscore = 8.2, RMSD = 3.9) and VirA of *Shigella* spp. (Zscore = 8.1, RMSD = 5.6). As for the comparisons within *Buchnera*, the aligned structure corresponds to the C-terminal region of SyeA (Fig 2b). Similarly, of the few SyeA homologs detectable by sequence-based searches, that of *Liberobacter* has been noted to have structural similarity to EspG^21^.

Both EspG and VirA are known T3SS effectors that interact with conserved eukaryotic innate immunity factors to facilitate invasion and persistence in host cells^22–24^, leading us to hypothesize that SyeA plays a related role in aphid bacteriocytes.

### SyeA predicted to interact with host Rho1

To investigate functions of SyeA, we explored whether it is predicted to interact with any host proteins. We focused on proteins involved in cytoskeletal rearrangement and lysosome trafficking, based on the known activities of EspG and VirA and on evidence that *Buchnera* is ultimately destroyed by lysosome fusion with symbiosomes^6^. Of 404 *A. pisum* proteins shown to be upregulated in bacteriocytes^25^, we chose 90 candidates, including 69 proteins annotated as being related to cytoskeletal rearrangement, cell trafficking, immunity, proteolysis, or signaling and an additional 21 proteins having unknown function. We obtained three predictions for interactions of each candidate with the C-terminal region of SyeA (amino acids 174-367) in AlphaFold3 (https://alphafoldserver.com/)^26^ and averaged the interface predicted template modeling (ipTM) score for each (Dataset S7). The protein with the highest average score was Rho1 (ipTM = 0.74, pTM = 0.80) (Fig 2c), a homolog to RhoA in humans and Rho1 of *Drosophila melanogaster*.

Rho family GTPases are central regulators of actin cytoskeletal rearrangement^27^. They are common targets of type III effectors secreted by bacterial pathogens^28,29^, with examples including VopS from *Vibrio parahaemolyticus*^30^ and YopE from *Yersinia pestis*^31^.

### Expression and detection of SyeA

We next sought to express SyeA heterologously in an *E. coli* system. We expressed both full-length SyeA and a truncated version (SyeAΔ) lacking the 172 amino acids of the N-terminus. In both cases, SyeA was detectable, but only within inclusion bodies, indicating protein aggregation and insolubility (Extended Data Fig 3a). Using a custom anti-SyeA polyclonal antibody, we demonstrated specificity through Western blots with controls consisting of *E. coli* harboring the same plasmid but lacking *syeA* (Extended Data Fig 3b-d). Using a cell-free expression system, we were able to express soluble SyeAΔ (Extended Data Fig 3e), indicating that insolubility is due to processes within cells.

Previously, *syeA* and *syeB* were shown to be actively transcribed and translated in *Buchnera* ^12,17,32^. We detected SyeA in samples from whole aphids and from bacteriocytes dissected from *A. pisum* individuals. In the former experiment, the full length SyeA (43.3 kDa) was detected in *A. pisum*, and in the closely related *A. kondoi* (Extended Data Fig 3e). In the experiment with dissected bacteriocytes, a truncated version (∼34 kDa) was detected (Extended Data Fig 3f). This smaller size approximates that expected if the first N-terminal alpha helix is lost, potentially during sample preparation method or post-translational processing within the bacteriocytes.

### SyeA peaks with colonization of embryos

To explore potential roles of SyeA in *Buchnera* colonization and persistence, we examined its expression at different host developmental stages, using immunofluorescence microscopy with anti-SyeA primary antibodies. Specificity was confirmed in both embryos and bacteriocytes through controls consisting of embryos prior to *Buchnera* colonization and controls lacking primary antibodies, imaged under the same conditions (Extended Data Figs 4a, 5a, b, e).

During asexual aphid reproduction, embryogenesis occurs within the mother’s abdomen, with each ovariole containing a succession of embryos at different developmental stages^9^. *Buchnera* cells are exocytosed from a nearby maternal bacteriocyte, losing the host-derived membrane, and endocytosed by the exposed syncytial bacteriome of blastoderm-stage (Stage 7) embryos, gaining the host-derived membrane which defines the symbiosome^5,8^. At Stage 10, the syncytium divides into uninucleate bacteriocytes which persist in mature aphids and which are degraded in older hosts as symbiosomes are destroyed through lysosomal fusion^6^.

The highest expression of SyeA occurs concurrently with colonization of the syncytial cytoplasm (Stages 7-8) (Fig 3, 4a, Extended Data Fig 5c-d). SyeA is elevated as soon as *Buchnera* is endocytosed, at the exposed edge of the syncytial membrane where the symbionts colonize (Fig 3c). As *Buchnera* moves through the syncytial cytoplasm, SyeA forms a ring at the periphery of each symbiosome (Fig 3, Fig 4a). Because embryos were dissected from mothers, it is not possible to establish whether elevated expression starts before entering the syncytium, before exocytosis from the maternal bacteriocyte or during the brief extracellular phase in the aphid body cavity.

**Fig 3.**
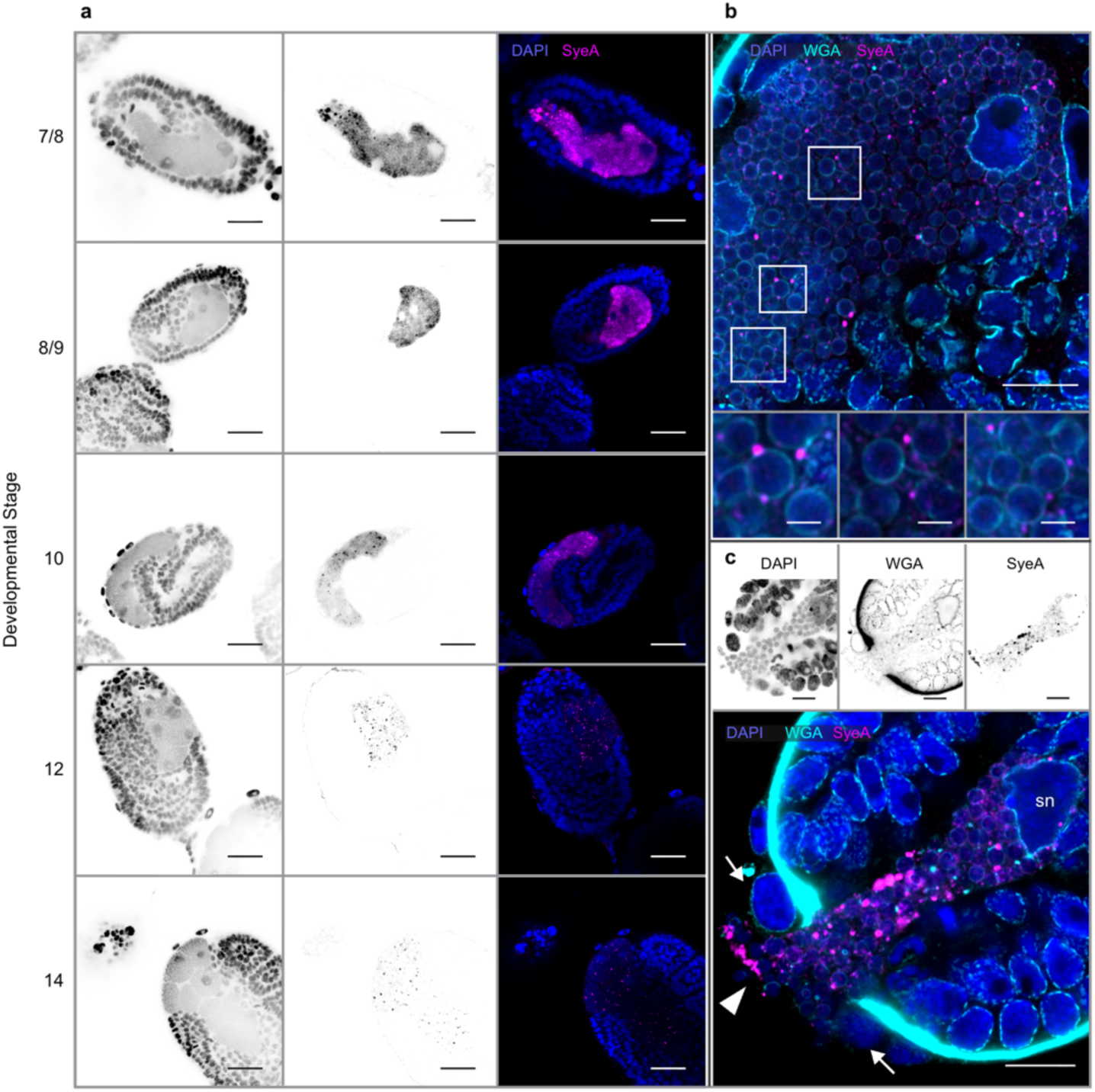
SyeA expression during early embryonic stages. **a**) Stage 7/8: SyeA abundant at entry point (arrowhead) and throughout syncytial bacteriome. Stage 8/9: Germ band invaginating, SyeA abundant in syncytium, Stage 10: Syncytium begins to form uninucleate bacteriocytes, accompanied by reduction in SyeA. Stage12: SyeA reduced in uninucleate bacteriocytes. Stage 14: Bacteriocytes pushed to the anterior and dorsal side, SyeA levels low. **b**) Stage 8 embryo showing location of SyeA exterior to symbiosome membrane. **c**) High SyeA expression immediately upon *Buchnera* entry into the syncytium, which extrudes from the embryo through the cuticle, flanked by large follicular (maternal) cells (arrows). In b) and c) WGA stains aphid membranes, *Buchnera* peptidoglycan, and chitin of exoskeleton. sn = syncytium nucleus. Scale bars: 20 μm in **a**, 10 μm in **b** & **c**, 2 μm in **b** expanded views. (**a**, **b**, and **c** represent separate experiments.)

The SyeA signal remains very strong until the syncytium divides into uninucleate bacteriocytes at Stage 10. At this stage, the signal diminishes sharply and is largely confined to small punctate clusters (Fig 4b-c, Extended Data Fig 5g) but is sometimes diffused across symbiosomal compartments (Fig 6b-c, Extended Data Fig 4d). This pattern continues in mature bacteriocytes (Fig 5d, Extended Data Fig 4 & 6).

**Fig. 4.**
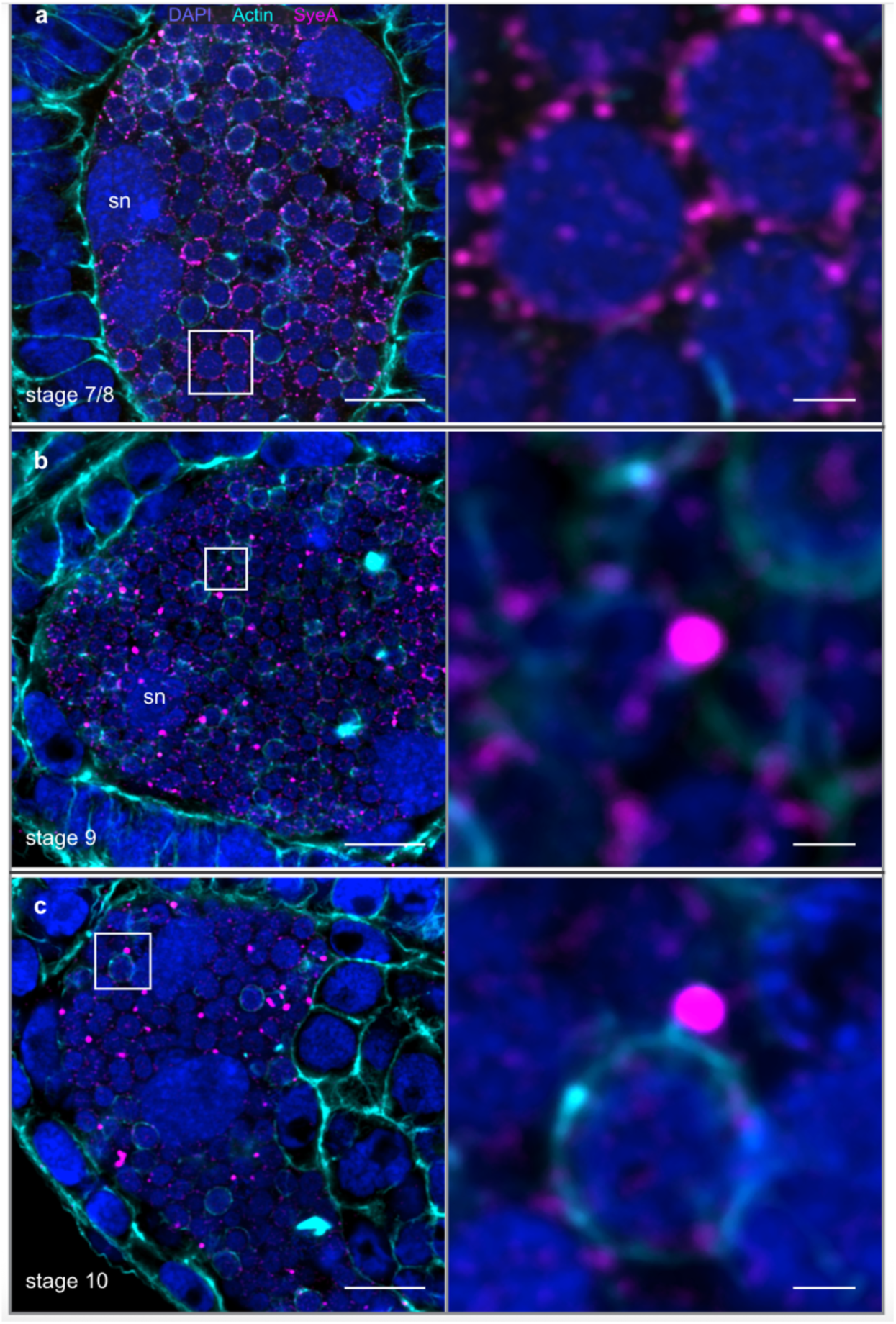
SyeA localization relative to that of actin during embryonic development. **a**) Stage 7/8: Immediately after colonization SyeA forms a dense ring at the border of the *Buchnera* cell but often appears to be within the symbiosomal membrane with a layer of actin outside. **b**, **c**) In Stage 9 and Stage 10, each symbiosome is surrounded by an actin layer, with SyeA located in bundles outside the actin, in the bacteriocyte cytosol. sn = syncytium nucleus. Scale bars: 10 μm in left column, 1 μm in right column.

**Fig. 5.**
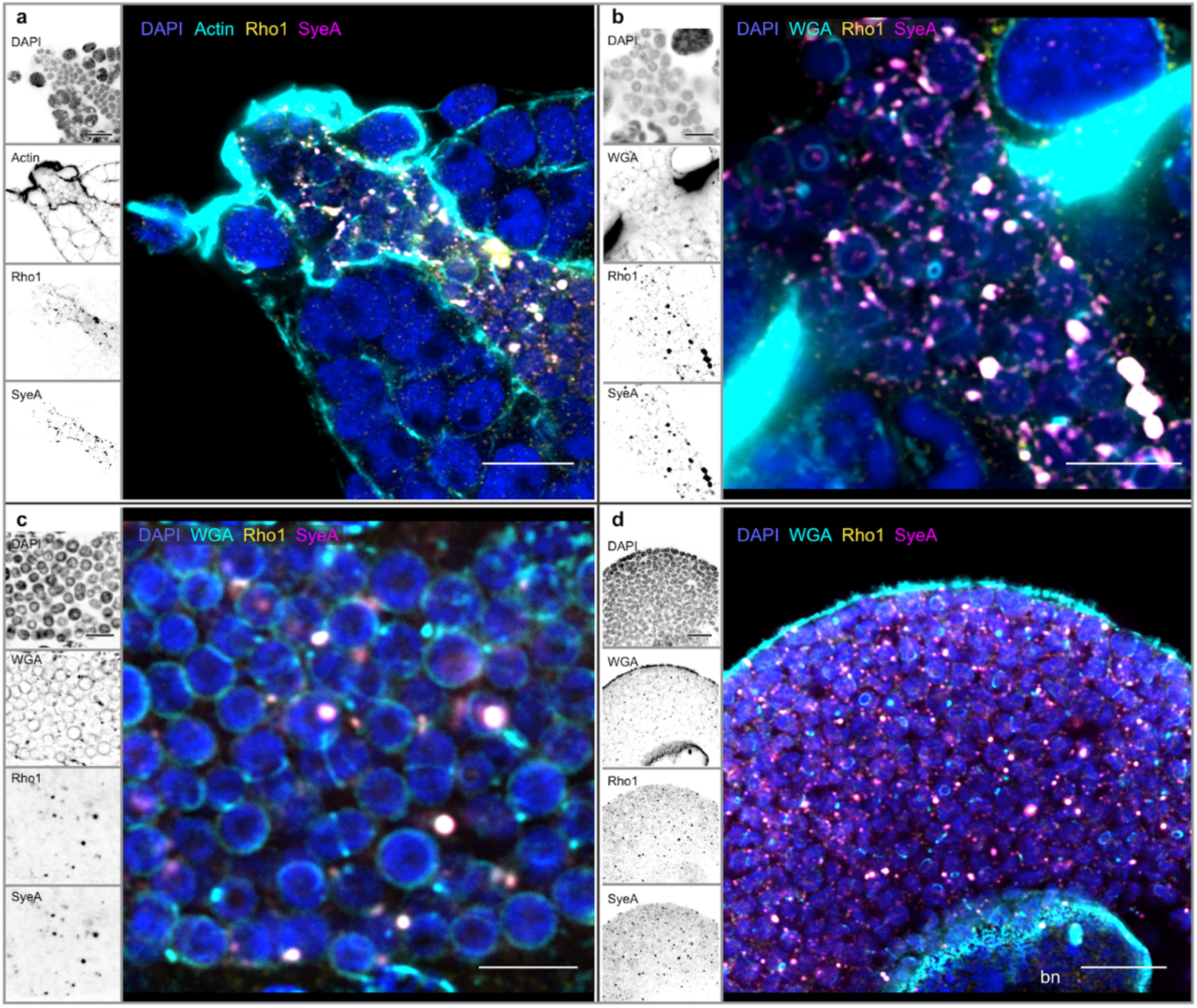
Colocalization of SyeA and Rho1 during colonization of embryo and in mature bacteriocyte. **a**) Actin mass at entry and along colonization channel surrounding syncytium in Stage 7 embryo. Arrows indicate enlarged, actin-rich follicle cells. **b**) Colonization channel in Stage 7 embryo showing colocalization of SyeA and Rho1 outside membranes. **c**) Closeup of syncytial bacteriome of stage 8 embryo, showing colocalization of SyeA and Rho1 in clusters outside symbiosomal membranes. **d**) Mature bacteriocyte of 8-day-old aphid showing reduced levels of both SyeA and Rho1, colocalized outside membranes surrounding symbiosomes. Cyan stain is phalloidin (actin) in **a**, WGA (membranes, chitin) in **b**-**d**. bn=bacteriocyte nucleus. Scale bars: 10 μm in **a** & **d**, 5 μm in **b** & **c**. (**a**-**d** represent separate experiments.)

### SyeA distribution in host cell cytoplasm

Using fluorescently labelled phalloidin, which stains actin, we found that the peak expression of SyeA at colonization coincides with appearance of a large mass of actin forming an extruding plug at the *Buchnera* entry point into the syncytium (Fig 5a, Extended Data Fig 7). This actin mass is reminiscent of the actin pedestals induced by enteropathogenic *E. coli*^33^.

Immediately following colonization of the syncytium, SyeA forms a layer at the symbiosome boundary, closely adjacent to, or overlapping, a layer of actin that surrounds the symbiosomal membrane (Fig 4a, Extended Data Fig 5c,d,g). As *Buchnera* moves down the syncytium, each symbiosome is surrounded by a layer of actin (Fig 4b-c). Later (Stage 10 and beyond), each symbiosome is surrounded by an actin layer, and most SyeA is outside this layer, in the bacteriocyte cytoplasm (Fig 4b-c). Thus, roughly coinciding with the formation of uninucleate bacteriocytes, SyeA is translocated across both the *Buchnera* cell wall and the symbiosomal membrane.

We examined mature bacteriocytes of aphids from 5 to 22 days old and found that SyeA is continuously detectable but at lower levels than in the embryonic syncytium and is usually confined to small clusters outside individual symbiosomes (Fig 5d, Extended Data Fig 4d, Extended Data Fig 6). Occasionally, SyeA is diffused across symbiosomal compartments (Fig 6b-c, Extended Data Fig 4d), as sometimes observed in symbiosomes near the nucleus (Fig 6c) where *Buchnera* degradation through lysosomal activity is initiated as the aphid ages^6^.

**Fig 6.**
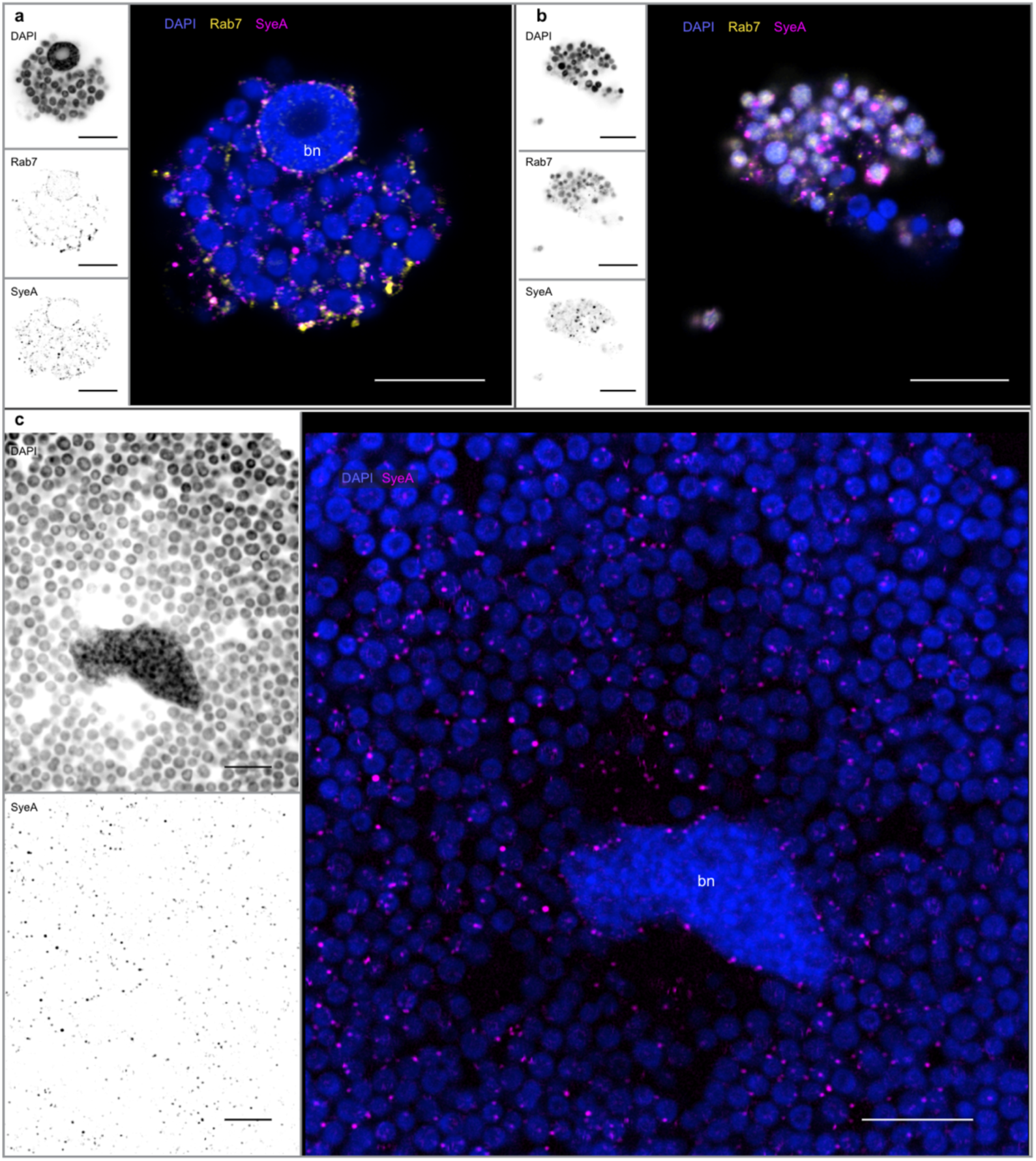
SyeA in mature bacteriocytes at different stages of degradation. **a**-**b**) Signals for SyeA and Rab7 are correlated, but not colocalized. SyeA increases and becomes diffuse in symbiosomal compartments in which *Buchnera* is being degraded, and Rab7 is most concentrated at the same compartments. **c**) SyeA distribution in bacteriocyte that is beginning to undergo degradation, with loss of *Buchnera* and diffuse SyeA (arrows) concentrated in symbiosomes near the bacteriocyte nucleus. bn=bacteriocyte nucleus. Scale bars: 10 μm. (**c** represents a separate experiment from **a** and **b**.)

### SyeA colocalizes with Rho1 of the host

EspG and VirA are known to interact with GTPases and to affect cytoskeletal rearrangements, and rho family GTPases are key regulators of the actin cytoskeleton^27^. Furthermore, our structural predictions suggest an interaction of SyeA with aphid Rho1 (Fig 2c). Therefore we examined the expression of Rho1 relative to that of SyeA, within embryos and bacteriocytes, using an anti-Rho1 antibody.

SyeA shows striking colocalization with Rho1 in both the embryonic syncytium and in bacteriocytes in both embryos and mature bacteriocytes (Fig 5, Extended Data Fig 6). These observations support a SyeA-Rho1 interaction in the bacteriocyte cytoplasm.

### SyeA and Rab7 abundances covary

Rab7 is a marker for late endosomes and has been shown to increase in aphid bacteriocytes in which *Buchnera* compartments are fusing with lysosomes and being degraded^6^. In mature bacteriocytes, we observed correlated expression of Rab7 and SyeA, with elevated levels of both proteins around certain symbiosomal compartments (Fig 6a-b). SyeA and Rab7 did not colocalize, indicating that SyeA does not directly interact with Rab7.

### *syeA* knockdown disrupts colonization

Because SyeA expression has structural homology to proteins enabling pathogenic bacteria to manipulate host cells and because SyeA levels peak at the time of colonization, we hypothesized that SyeA is required for colonization. We predicted that disrupting its function would interfere with colonization and disrupt embryonic development, which does not proceed normally in the absence of *Buchnera*^7^.

Although genetic manipulation of intracellular symbionts including *Buchnera* has not yet been possible, recent studies have used peptide nucleic acids (PNAs) to reduce expression of targeted genes in *Buchnera*-Ap, resulting in phenotypic changes affecting symbionts and hosts^34,35^. We applied this method to observe effects of *syeA* knockdown on the symbiosis. Based on the potential roles and observed expression of SyeA, we examined both embryos and mature bacteriocytes for phenotypes related to its knockdown (Fig 7a).

**Fig 7.**
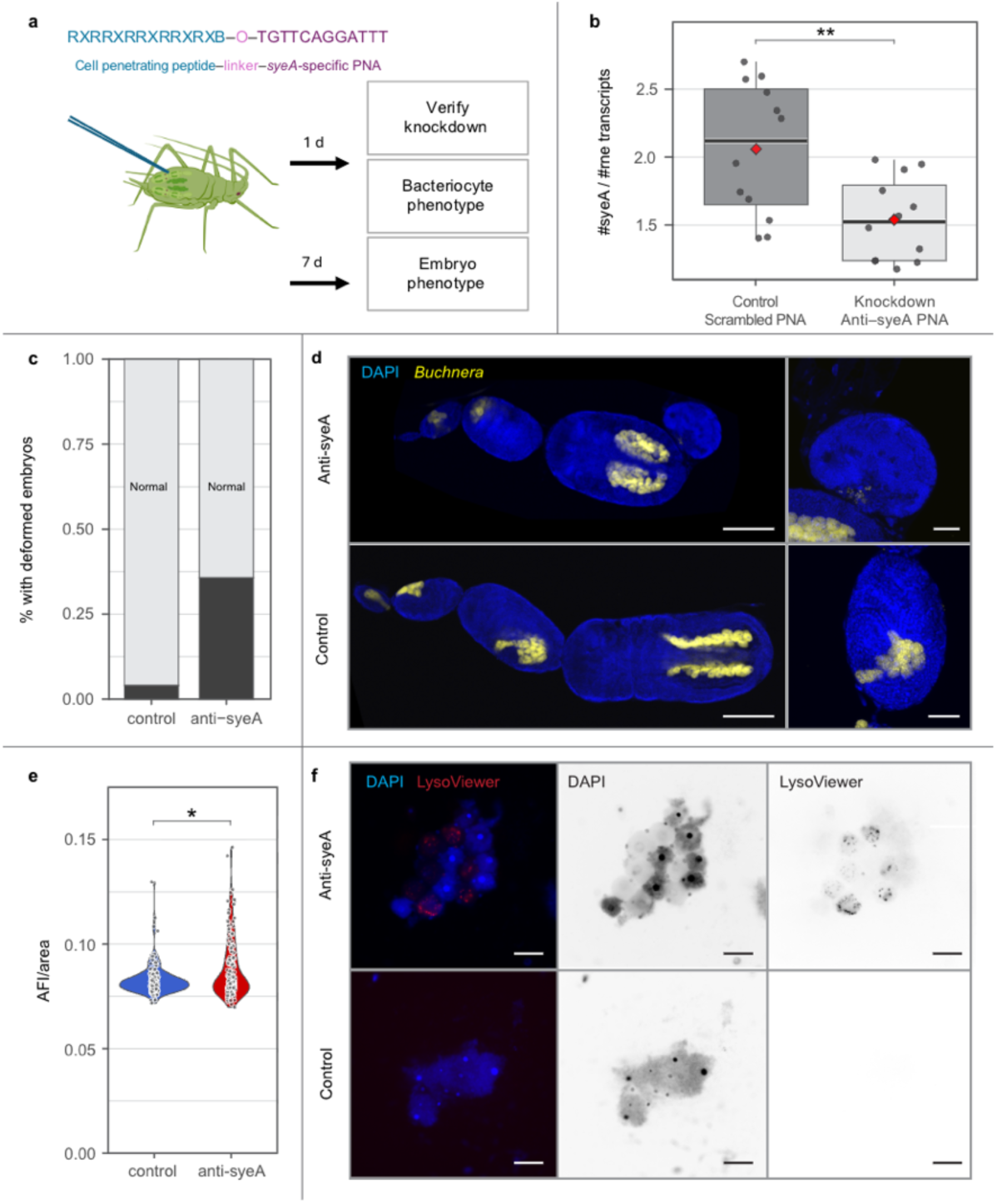
Effects of *syeA* knockdown on colonization and development of host. **a**) anti-*syeA* PNA and experimental design. **b**) Validation of *syeA* knockdown by dPCR of transcript levels relative to those of control gene *rne.* **c**) Percent of mothers with deformed embryos for control and treated aphids. **d)** Representative images of late-stage phenotypes of embryos that were at stage 7 at time of exposure (images collected 7 days after injection of PNA). Top panel shows ovariole with deformed and stunted embryo in terminal position in anti-syeA PNA treatment with a closeup of the affected embryo, lower panel shows normal terminal embryo of control treatment and shows a normal embryo of the same size but earlier developmental stage. **e**) Effect of *syeA* knockdown on lysosomal degradation of *Buchnera* 1 day after treatment. Graph shows fluorescence (AFI) per area per bacteriocyte, as detected using LysoViewer. Each point represents one bacteriocyte. **f)** Representative images of LysoViewer signal for treated and control aphids. (Aphid graphic: M. Steele.) Scale bars: 200 μm in left hand images of **d**, 5 μm in right hand images of **d**, 100 μm in **f**.

We designed an anti-*syeA* PNA and a control PNA and injected it into 4-day-old aphids (Methods). To verify knockdown success, we sampled 24 h later and measured *syeA* transcript levels by making cDNA and performing digital PCR. We found significantly less *syeA* transcript in the anti-*syeA*-treated aphids, with levels averaging 49% of levels in control aphids (Wilcoxon rank sum test with continuity correction: W = 122, *p* = 0.0043) (Fig 7b, Dataset S8).

To determine if *syeA* knockdown causes defects in colonization or embryonic development, 7-day-old (4th instar) aphids were injected with control or anti-*syeA* PNAs and their embryos examined 7 days later. This timing enables us to examine how knockdown at the colonization stage (Stage 7) impacts embryos, which would be in terminal positions (∼Stage 19, approaching birth) at our sampling time^9,36^.

Many terminal embryos dissected from aphids treated with anti-*syeA* PNA exhibited pronounced deformities and stunted growth, whereas embryos of mothers treated with the control PNA were normal (Fig 7c, Supplementary Movie 2) (*p* = 0.0058, Fisher’s exact test, N = 25 control, 28 treated). Thus, SyeA appears to be essential for successful colonization of embryos and consequently for successful development.

### *syeA* knockdown boosts lysosome activity

To examine the effects of *syeA* knockdown on maternal bacteriocytes, 4–5 day-old (3rd instar) aphids were injected with control PNAs or anti-*syeA* PNAs and dissected ∼24 h later. Live cell imaging was performed using an acidotropic dye (LysoViewer640) marking lysosomal compartments. Quantification of average fluorescence intensity per unit area showed that anti-*syeA*-treated aphids exhibited elevated lysosomal activity for symbiosomal compartments (*p* = 0.0323, Mann-Whitney U) (Fig 7d). This result supports a potential role of SyeA in suppressing lysosomal destruction of *Buchnera* within aphid bacteriocytes.

## DISCUSSION

The first *Buchnera* genome sequence^3^ enabled insights into the evolution of this symbiont and its role in hosts, but also presented new puzzles. In the face of extreme gene loss, the retention of highly expressed flagellar genes^12,13,15^ implied an important function, suggesting that *Buchnera* secretes other protein(s) of unknown identity^3^. Also noted in the original genome publication was the presence of the orphan genes *syeAB* (originally *yba34*), for which functions have remained entirely unknown.

Our results show that SyeA is secreted outside the *Buchnera* cell and translocated across the symbiosomal membrane, into the bacteriocyte cytoplasm, where evidence supports an interaction with Rho1. Despite rapid sequence evolution, SyeA retains structural similarity across distant *Buchnera* lineages and has structural homology to effectors of enteric pathogens. Further, SyeA is highly expressed upon transmission to embryos, is required for colonization, and appears to suppress lysosomal destruction in mature bacteriocytes.

In pathogenic *E. coli* and *Shigella*, the T3SS effectors EspG/VirA^22,23^ contribute to evasion of cell-intrinsic immune responses^37,38^. They target GTPases and thereby affect host cytoskeletal structure and lysosomal and endocytic pathways^39^. VirA is essential for *Shigella*’s virulence and is involved in cell-cell spread, host cytoskeletal rearrangement, and escape from lysosomes^40^. Although EspG and VirA sequences are divergent (∼20% AA identity), EspG can rescue *Shigella* virulence in null VirA mutants^22^. In *Shigella,* VirA is required to colonize host cells via the remodeling of actin scaffolding and consequent ruffling of the host membrane^23^, a process dependent on Rho family proteins^41^. In *Buchnera*, each symbiosome is surrounded by an actin sheath (Fig 4b-c), a pattern reminiscent of the actin around *Shigella-*containing vacuoles following entry into host cells^42^.

In animals generally, Rho family proteins regulate the actin cytoskeleton^27^. *Buchnera* colonization of the syncytial bacteriome coincides with actin buildup near the entrypoint (Fig 5a, Extended Data Fig 7), as observed previously^8,9^. Thereafter, each symbiosomal compartment is surrounded by actin, both in the syncytium and in mature bacteriocytes. Our images suggest direct interaction between SyeA and Rho1 (Fig 5, Extended Data Fig 6), which colocalize throughout development. Both are elevated during colonization, up to the stage at which uninucleate bacteriocytes form. Because we examined embryos dissected from mothers, we could not determine whether SyeA expression is prior to colonization, in nearby maternal bacteriocytes or in the extracellular transmission phase.

Another apparent function of SyeA is suppression of lysosomal activity within symbiosomal compartments in mature bacteriocytes. This hypothesis is supported by the observation that SyeA abundance increases in bacteriocytes undergoing degradation, and that knockdown of *syeA* results in increased number of symbiosomes with lysosomal activity (Fig 7d). Rab7, a marker for late endosomes destined for degradation, shows increased expression around such symbiosomes^6^, and SyeA expression correlates with that of Rab7 in mature bacteriocytes (Fig 6a-b).

The localization of SyeA in the bacteriocyte cytoplasm implies that it is not only secreted from the *Buchnera* cell but also translocated across the symbiosomal membrane. How translocation happens remains unknown: flagellar basal bodies are not known to function as injectisomes as do canonical T3SSs^11^. Other mechanisms for translocation, such as an accompanying translocator protein, are possible^43^. Potentially movement across the host membrane is dependent on the variable N-terminal region and/or SyeB as both have alpha helices along with disordered regions (Extended Data Fig. 1), and SyeB is predicted as a Type 3 effector (Extended Data Table 1). In this mutualistic symbiosis, the host is also expected to evolve mechanisms to support the symbiosis, potentially including adaptations facilitating the translocation^44^.

EspG and VirA, along with other pathogenicity factors, are encoded within horizontally transferred genomic islands that encode numerous proteins enabling pathogenicity^22–24^. We hypothesize that a free-living *Buchnera* ancestor acquired a genomic island containing *syeA* and that this acquisition enabled an intracellular phase. Initially invasion was likely strictly bacterium-dependent, with SyeA, and possibly other effectors, enabling endocytosis and suppression of the lysosomal response of hosts, as observed for T3SS effectors of pathogens.

Following the transition to a host-beneficial state, selection would favor host modifications that ensure transmission and control of the endosymbiont. For example, aphids have acquired bacterial genes needed for *Buchnera*’s peptidoglycan production^45^. In aphids and many other symbioses, such adaptations have led to orderly incorporation into host development and confinement of the symbionts to specialized cells. Increased host control could lead to erosion of the original bacterial invasion mechanisms. Our results imply that SyeA remains essential for the *A. pisum*-*Buchnera* symbiosis, perhaps as a vestige of invasion mechanisms that were originally more elaborate. Several other *Buchnera* lineages appear to have progressed further in this erosion, having lost SyeA, flagellar genes (Fig 1a-b), and machinery for making most cell wall components^46^.

*Buchnera* falls within a bacterial clade that includes many obligate endosymbionts of insects, with varying levels of genome reduction^1,47,48^. Several more recently evolved endosymbionts maintain intact T3SSs, which can be essential for colonization^49^. More ancient symbionts, including *Buchnera*, have smaller genomes and appear to be largely under host control. Our findings for *Buchnera* suggest that such endosymbionts can originate as pathogens that invade hosts using T3SSs or other secretion systems.

## Acknowledgements

We thank Derrick Kamp for help with microscopy images and feedback, Katherine Tan for advice on PNAs, Kim Hammond for maintenance of aphid colonies and figure preparation, Michelle Mikesh, Anna Webb and Quinn Lee for help with microscopy, and Howard Ochman for feedback. Microscopy and proteomics were performed at the UT Austin Center for Biomedical Research Support Microscopy and Flow Cytometry Facility (RRID:SCR_021756) and Biological Mass Spectrometry Facility (RRID:SCR_021728). Funding was from National Institutes of Health awards R35GM131738 and R35GM158145 to NAM. **Data Availability**: Mass spectrometry proteomics data were deposited to the ProteomeXchange Consortium via the PRIDE partner repository under dataset identifier PXD066113

## Code availability

This study did not generate any new code.

## Author affiliations

All authors performed this work while affiliated with the Department of Integrative Biology, The University of Texas at Austin, Austin, TX, USA

## Author contributions

GPM, AZX, EWY, AS, JEP, and TES performed laboratory experiments. GPM and AZX performed bioinformatic analyses. GPM and NAM originated and directed the project and wrote a draft of the manuscript. All authors contributed to experimental design and reviewing the manuscript. NAM provided funding and facilities.

## Competing interests

The authors declare that they have no competing interests.

**Extended Data** are available for this paper.

**Supplementary Information** is available for this paper. Correspondence and requests for materials should be addressed to.

Other supporting materials for this manuscript include the following: Datasets S1-S8 (in a single Excel file)

Extended Data Movie 1: Structural alignment of SyeA from divergent *Buchnera*

Available at: https://drive.google.com/file/d/1413XZR6EojLDDNvuqccptPK-T2oS_2SD/view?usp=drive_link

Extended Data Movie 2. Embryos from PNA experiment.

Available at: https://drive.google.com/file/d/1DILjWMiLiRu0sFH41BG3X7lLqMnKcW43/view?usp=drive_link

**Legends for Datasets S1 to S8**

**Dataset S1.** Predictions as Type 3 Secretion System effectors for all protein sequences of the *Buchnera* APS genome (NCBI NC_002528), as calculated using BastionX.

**Dataset S2.** List of *Buchnera* genomes used for OrthoFinder analysis. Aphid subfamily classification derived from Jousselin et al. 2024. Tribe classification is included only for members of Aphidinae.

**Dataset S3.** List of OrthoGroups used for presence/absence analysis presented in Fig 1a. Protein names and accession numbers are taken from the Buchnera APS genome (NCBI NC_002528).

**Dataset S4.** Presence and absence of *syeA* and *syeB* and percentage of flagellar genes for all 115 *Buchnera* genomes analyzed. The first three columns correspond to the results presented in Fig 1a.

**Dataset S5.** List of 103 *syeA* orthologs, positions within *Buchnera* genomes, and predictions as Type 3 effectors by BastionX.

**Dataset S6.** Presence and absence of 26 flagellar genes in all 115 *Buchnera* genomes.

**Dataset S7.** Prediction for interactions with C-terminal region of SyeA of *Buchnera-*Ap for pea aphid genes, based on 3 prediction runs with AlphaFold3.

**Dataset S8.** Results from dPCR validation of PNA knockdown of *syeA.* These data correspond to results in Fig 7b.

## Methods

### Computational prediction of T3SS effectors

To determine which *Buchnera* proteins are most likely to be secreted by a T3SS, the complete proteome of *Buchnera*-Ap (Uniprot ID UP000001806) was run in BastionX, an online tool that predicts the likelihood of a given protein to be a substrate for bacterial secretion systems (https://bastionx.erc.monash.edu/server.jsp)^1^. The benchmarking parameter for v2.0 of BastionX was selected for the run (Dataset S1). An earlier version, Bastion3 had a false positive rate of 4.1%^1^, and this rate is expected to be lower in BastionX.

To determine the likelihood of SyeA orthologs from diverse *Buchnera* being substrates for a T3SS, a total of 103 SyeA orthologs were run in BastionX, with the common use parameter for v2.0 selected (Dataset S2).

### *syeA*, *syeB*, and flagellar genes in *Buchnera*

To survey the presence and absence of *syeA*, *syeB*, and flagellar basal body genes in a wide range of *Buchnera* strains, genomes of *Buchnera* strains uploaded to NCBI GenBank were selected for comparative orthology analysis (Dataset S2). The most recently uploaded *Buchnera* genome from each available host aphid species (April 22, 2025) was chosen. Contig- and scaffold-level assemblies were excluded. If available, the RefSeq annotation was used over the GenBank annotation. A total of 115 proteomes were collected and run on OrthoFinder v3.0 (default parameters) (Emms and Kelly 2019) to generate orthogroups. Orthogroups corresponding to the 26 flagellar basal body genes present in *Buchnera-*Ap (12 *flg*, 2 *flh* genes, and 12 *fli* genes), along with *syeA* and *syeB*, were identified using the known accession numbers of the protein sequences from *Buchnera*-Ap (Dataset S3). Percentage of total flagellar basal body genes were calculated for each species (Dataset S4).

Genomes deemed by OrthoFinder to lack *syeA* were manually examined to check for the presence of divergent *syeA* homologs. Genomic rearrangements are rare in *Buchnera*^2^. To examine the gene neighborhood across *Buchnera* strains, we selected conserved neighboring genes (*hslV*, *hlsU*, and *rho*) retained in all lineages and flanking *syeA* (Fig 1b). We downloaded these regions as fasta files from NCBI, re-annotated them with prokka (v1.14.6)^3^ and used Clinker^4^ and GeneGraphics^5^ for calculating pairwise sequence similarity and for visualization. If present, the corresponding protein sequence was run through NCBI BLASTp to check for hits against SyeA orthologs in other *Buchnera* strains. In two cases (*Buchnera* of *Cavariella theobaldi* and *Pterocomma populeum*), the *syeA* region contained a short hypothetical protein on the strand normally encoding *syeA* flanked by expanded stretches of non-coding DNA, but the hypothetical proteins showed no detectable similarity to SyeA, which was thus scored as missing. Likewise, *Buchnera* of tribe Macrosiphini encoded SyeB with recognizable sequence homology across species, whereas *Buchnera* of tribe Aphidini encoded a short protein in the same position, but without sequence similarity to SyeB.

To visualize presence and absence patterns of *syeA*, *syeB*, and flagellar genes across *Buchnera*, a published *Buchnera* phylogeny^6^ was used to create a cladogram using a select number of *Buchnera* genomes from host aphid species representing all major subfamilies (Fig 1a). Less-supported nodes were collapsed.

### Rate of SyeA sequence evolution

To examine rates of evolution of *Buchnera* proteins, coding sequences (CDS) for *Buchnera* strains APS (GCF_000009605.1), *Acyrthosiphon kondoi* (GCF_000225445), and *Schizaphis graminum* (GCF_000007365.1) were accessed from NCBI. Calculations were implemented in R (version 4.3.0), using the package Orthologr^7^. Only orthologous genes shared by all three strains were included. The dNdS module was used to estimate *d*N (number of nonsynonymous substitutions per nonsynonymous site) for each gene of *Buchnera*-Ap paired with its ortholog in each of the other two species.

### Structural similarity searches

To examine structural similarity across divergent *Buchnera* strains, AlphaFold-predicted structures were used for comparison (AlphaFold Monomer v2.0). Previously computed .pdb files were downloaded from Uniprot, and similarity was assessed using FoldMason on the FoldSeek online server^8^. Similarity was also scored using the DALI protein server^9^.

For similarity searches in genomes other than *Buchnera*, SyeA of *Buchnera*-Ap (UniProt P57644) was used as a search query against the PDB. Additionally, structural searches were performed against the AlphaFold-predicted protein database AFDB. To assess structural alignment quality, pairwise comparisons between SyeA of *Buchnera*-Ap (UniProt ID P57644), EspG, and VirA (UniProt IDs Q5WMC0 and Q7BU60) were performed using DALI and the resulting Z-scores and RMSD values were obtained. We also downloaded AlphaFold-predicted structures for the conserved C-terminal region of SyeA for several distantly related *Buchnera* lineages (*Buchnera*-Cc, *Buchnera*-Ua, *Buchnera*-Bp, corresponding to UniProt IDs Q5EU88, G2LQ59, Q89A26) and compared these to that for *Buchnera*-Ap to assess extent of structural similarity across *Buchnera*.

### Antibody production

Two Anti-SyeA polyclonal antibodies against SyeA of *Buchnera*-Ap were generated by a third-party service (BioMatik, Ontario, CA) by generating reactive sera in rabbit against a conserved SyeA peptide (amino acids 272-286) and against a heterologously expressed full-length His6-tagged SyeA. The secondary antibody was goat anti-rabbit-AlexaFluor647 (ThermoFisher A21245).

The Rho family and Rab7 GTPases are highly conserved, allowing us to use available antibodies for other animal species. For visualizing Rab7, we used a mouse IgG2b monoclonal antibody (Sigma-Aldrich R8779), previously shown to have specificity for Rab7 in *A. pisum*^10^. The secondary antibody was goat anti-mouse IgG2b-AlexaFluor568 (ThermoFisher A21144). The primary antibody for Rho1 was mouse IgG1 raised against full-length Rho1 of *Drosophila melanogaster* (https://dshb.biology.uiowa.edu/ p1D9). Rho1 proteins from *D. melanogaster* (UniProt P48148) and *A. pisum* (UniProt ACYPI003261) contain 192 amino acids and are 94% identical. The secondary was goat anti-mouse IgG1-AlexaFluor568 (ThermoFisher A21124). Rho1 and Rab7 were visualized in separate experiments.

### Heterologous expression of SyeA

To confirm the specificity of the anti-SyeA antibody, we heterologously expressed His6-tagged SyeA. We used *E. coli* Arctic Express, a BL21 expression strain that is compatible with inducible expression plasmids and that contains plasmids expressing chaperone proteins from the cold-adapted bacterium *Oleispira antarctica*^11^. Use of molecular chaperones and low induction temperatures can facilitate expression of thermally unstable endosymbiont proteins^12^.

The *syeA* gene was amplified from *Buchnera* genomic DNA purified from a lab colony of *A. pisum* strain LSR1 and cloned into a pET28b vector which added a His6-tag and placed it under an inducible T7 promoter. The resulting plasmid was transformed into Arctic Express via electroporation. A control was made using the same plasmid but lacking the *syeA* region. To determine suitable expression conditions for SyeA, we used overnight Arctic Express cultures to inoculate fresh LB cultures at a starting OD of 0.1 and grew them at 30℃ until they reached exponential phase (OD∼0.4-0.6), when they were either kept at 30℃ or transferred to 15℃ or 4℃. Protein expression was induced by the addition of various concentrations of IPTG, ranging from 0.1 mM to 1 mM. Cultures were grown for up to 24 h post-induction.

To detect expressed SyeA, culture samples were pelleted and processed with bacterial lysis buffer and lysozyme to obtain soluble protein fractions. The remaining insoluble pellets were denatured and solubilized in urea and SDS. Both soluble and insoluble fractions of each temperature and IPTG concentration condition were analyzed by Coomassie staining or Western blot with anti-His6 tag antibodies. SyeA bands on both Coomassie staining and Western blot were seen only in samples of cultures grown at 30℃ and induced at 0.1 and 0.5 mM, with the best expression at 0.1 mM. However, the SyeA was found only in the insoluble fraction, indicating that SyeA was likely misfolding or aggregating in inclusion bodies (Extended Data Fig 3a). In further experiments, SyeA expression was improved using 0.05 mM IPTG at 30℃ (Extended Data Fig 3b-c).

We hypothesized that the non-conserved N-terminus region (Extended Data Fig 1, top left) might be causing aggregation and insolubility of SyeA in *E. coli*. Since only the C-terminus portion is conserved in *Buchnera*, we reasoned that further studies of SyeA may not be affected by and may even be helped by only expressing only this portion. Therefore, using a third-party service (Twist Bioscience, San Francisco, CA), we generated a gene fragment for producing SyeAΔ, a truncation construct lacking the 172 amino acids of the N-terminus, with an N-terminal His6 tag and codon-optimized for *E. coli* expression. PCR was used to add overhangs to the gene fragment, allowing for cloning into the pET28b vector using Gibson Assembly® Master Mix (New England Biolabs). The gene fragment was cloned into the same pET28b vector used for full *syeA* sequence and expressed using conditions most successful for the full-length SyeA (uninduced or induced at 0.05 mM IPTG at 30℃, sampled 24 hr post-induction). Cultures were pelleted, and pellets solubilized in 6X SDS dye, followed by running on a 12% acrylamide gel and Coomassie staining and Western blots. SyeAΔ showed better expression than full-length SyeA but remained in the insoluble fraction (Extended Data Fig 3d).

### Cell-free expression of SyeA**Δ**

As SyeAΔ was insoluble in expression experiments in *E. coli*, we used the NEBExpress® Cell-free *E. coli* Protein Synthesis System and added the pET28b-SyeAΔ construct. After 2-3 h of incubation at 37℃, we confirmed that soluble SyeAΔ was expressed by comparing protein bands to a negative control lacking added DNA (not shown). We purified SyeAΔ from pooled cell-free expression mixtures using a nickel resin spin column, yielding 0.5 mg/mL in the final eluate.

SyeAΔ was also expressed via the PURExpress® In Vitro Protein Synthesis Kit (New England Biolabs), using 0.25 μL of the vector and incubating at 37℃ for 2 h. Purification was confirmed with SDS-PAGE and Coomassie staining (Extended Data Fig 3e).

### SyeA detection in aphid bacteriocytes

To detect SyeA from aphid bacteriocytes, aphid samples of four species (*A. pisum*, *A. kondoi*, *M. persicae*, *Aphis craccivora*) were frozen at -80℃, thawed, flash-frozen in liquid N_2_, crushed to powder, suspended in 2X SDS loading buffer, homogenized by passing through a fine needle, diluted in 1X PBS, and boiled at 100℃ for 15 minutes, then run on SDS-PAGE. Western blotting was performed with anti-SyeA raised against a peptide. We also detected SyeA in bacteriocytes dissected from 100 14-day-old *A. pisum* str. LSR1 individuals. Following dissection in cold Buffer A (25 mM KCl, 35 mM Tris, 10 mM MgCl2, 250 mM sucrose, 250 mM EDTA, pH=7.5), bacteriocytes were homogenized in cell lysis buffer (25 mM Tris, 0.15 M NaCl, 1 mM EDTA, 1% Triton X-100, 5% glycerol), then centrifuged at 12,000 rpm for 20 min at 4℃ to remove cell debris. Westerns were performed using the anti-SyeA raised against full-length protein. For negative controls, samples containing only embryos and surrounding follicle cells from stages prior to *Buchnera* colonization were collected from five aphids.

To detect SyeA and SyeB with mass spectrometry, protein lysate samples from 7-day-old aphids were mixed with 6X SDS dye (0.6 M DTT, 0.35 M Tris pH 6.8, 30% v/v glycerol, 10% w/v SDS, 0.012% w/v bromophenol blue), boiled at 100℃ for 10 min, and run on a 12% acrylamide gel for 2 min. The gel was stained with Coomassie Brilliant Blue G-250 (3 g/L in 45% H_2_O, 45% methanol, 10% glacial acetic acid) for 10 min, then destained overnight in destain solution (85% H_2_O, 10% glacial acetic acid, 5% ethanol). The protein band was excised and run in LC-MS/MS shotgun proteomics by the University of Texas Biological Mass Spectrometry Facility. Two samples were run and analyzed using Scaffold5 (https://www.proteomesoftware.com). We detected total peptide spectra counts mapping to *Buchnera-*Ap of 112,094 and 109,193 in the two samples. Of these SyeA was represented by 6 and 6 spectra, and SyeB by 2 and 3 spectra in the two samples. Mass spectrometry proteomics data were deposited to the ProteomeXchange Consortium via the PRIDE^13^ partner repository under dataset identifier PXD066113.

To detect SyeA by Western blotting, protein lysate samples were mixed with 6X SDS dye, boiled at 100℃ for 10 min, and run on a 12% acrylamide gel at constant 120 V for 1 hr. Protein on the gel was subsequently transferred to a PVDF membrane using the Mini Blot Module (ThermoFisher Scientific) at 20 V and 400 mA for 1 hr. Membranes were washed in blocking milk buffer (5% skim milk powder in 1X TBST) for 1 hr at room temperature, then incubated with 10 mL primary antibody milk (1:10,000 anti-SyeA antibody A in blocking milk buffer) overnight at 4℃. Then, membranes were washed for 5 min twice with blocking milk buffer, then incubated with 10 mL secondary antibody milk (1:1,500 goat anti-rabbit AlexaFluor680 (Abcam, Cambridge, UK)) for 90 min at room temperature. Finally, membranes were washed three times for 5 min with blocking milk buffer and once for 10 min with 1X TBST, and imaged with the Odyssey CLx (LI-COR Biosciences, Lincoln, NE) at the 700 nm channel. To avoid potential loss of SyeA during the protein purification process, 50 μL of 6X SDS dye was added directly to bacteriocytes dissected from 100 LSR1 aphids and boiled at 100℃ for 10 min. 15 μL was loaded onto an SDS-PAGE gel, and a Western blot was performed using the anti-SyeA antibody at the same concentration as before (Extended Data Fig 3f).

To rule out cross-binding of secondary antibodies as a possible artifact underlying the observed colocalization of Rho1 and SyeA, we performed a Western in which the SyeA secondary was used against both primary antibodies (Extended Data Fig 3g).

### Immunofluorescence microscopy

Immunofluorescence microscopy followed a published protocol^10^. Antibodies described above were used to visualize SyeA, Rho1, and Rab7. Actin was visualized with phalloidin-CF488 (Biotium 0042T). Aphid membranes, peptidoglycan of the *Buchnera* cell wall, and chitin were visualized with WGA-CF488 (Biotium 29022). (Because *Buchnera* has lost outer membrane components, WGA is not expected to bind to its membrane^14,15^). DAPI was used to stain DNA.

For each experiment, same-aged aphids (age depending on experiment) were dissected in cold PBS to obtain embryos or bacteriocytes, and these were fixed in 3.7% formaldehyde in PBS on ice for 20–25 min. Samples were washed once in PBS and permeabilized in PAXD-Tween 20 (PBS, 5% BSA, 0.3% sodium deoxycholate, 0.3% Tween 20). The primary antibody (anti-SyeA, anti-Rho, or antiRab7) was diluted to 5 µg/mL in PAXD-Tween 20 and incubated overnight at 4°C. Samples were washed once in PBST and blocked with PAXD-Tween 20 for 30 min. Secondary antibody incubation was performed using a 1:200 dilution to 10 µg/mL in PAXD-Tween 20 with DAPI, and incubated at 4°C overnight. Samples were then washed with PBS twice and mounted in SlowFade Mounting Medium. Mounted samples were imaged using either a Nikon Eclipse epifluorescence microscope or a Nikon AXR-NSPARC laser scanning confocal microscope. Off-target binding was assessed with negative controls lacking primary antibody. Control and experimental samples were otherwise treated the same and imaged under the same settings.

### *syeA* Knockdown with peptide nucleic acid

Although genetic manipulation of *Buchnera* has not yet been possible, recent studies showed the successful application of peptide nucleic acids (PNAs) to reduce expression of targeted genes in *Buchnera*-Ap^14,16^. PNAs target the mRNA and interfere with translation or induce degradation of the transcript. A synthetic oligonucleotide complementary to the target gene is attached to a cell penetrating peptide, which has been shown to increase uptake by cells. We applied this method to observe effects of reducing expression of *syeA* on the symbiosis. We designed PNAs using MASON, a package for designing targeted PNAs with no off-target hits within a genome^17^. For the anti-*syeA* PNA, we used a sequence (TGTTCAGG<u>AAT</u>T) that targets the beginning of the coding sequence of *syeA* including the start codon (which is ATT for this gene, as AT-rich genomes such as *Buchnera* often use this alternative start codon). The control PNA used an oligonucleotide (AAGTCTTATGGT) of the same composition and length, but scrambled and designed to prevent off-target matches. The PNAs consisted of these oligonucleotides each linked to an arginine-rich cell-penetrating peptide (RXRRXRRXRRXRB-O-) to enable entry of the molecule into cells. PNAs were synthesized by PNA Bio Inc (Thousand Oaks, CA). Lyophilized PNAs were resuspended to 200 µM in water. Aphids were injected with approximately 0.1 µL of 100 µM PNA in 6 mM CaCl_2_.

### Validation of *syeA* knockdown

To verify that the anti-*syeA* PNA reduces *syeA* transcripts, RNA was extracted from single, whole aphids 24 h after injection, with 12 anti-*syeA* and 12 control individuals. Each aphid was transferred to a 1.5 mL pestle tube to which 200 μL of Tri Reagent (MRI, Cincinnati, OH, USA) was added, homogenized for 30 s with a plastic pestle (MilliporeSigma, St. Louis, MO, USA) to promote cell lysis, and kept on ice. RNA extraction was performed using the Zymo Direct-zol kit (Zymo Research, Irvine, CA, USA; product number R2052) following the protocol described in the kit including a DNase treatment and eluting with 50 μL molecular grade water. Concentration and integrity were checked using both Nanodrop (1 μL) and Qubit BR RNA (5 μL), and by running on an agarose gel. cDNA synthesis was performed using the PrimeScript cDNA synthesis kit (Takara Bio USA, San Jose CA, USA; product number RR037A) following the kit protocol and using 200 ng of total RNA (calculated based on Qubit concentrations) for each 10 μL reaction.

Digital PCR (dPCR) was performed on 2 μL of a 1/10 solution of the cDNA for each sample. Primers were: SyeA_4F (GAAGCGACTATTTCTTTATCTGAAC) and SyeA_4R (AGGATGCCGTGGATTATTATTT) and were first tested with serial dilutions of aphid cDNA for low variability and linearity of reported product concentration. Assays were run on a Qiagen QIAcuity One, 2-plex digital PCR machine (Qiagen,Germantown, MD, USA) with a 24-well 26K nanoplate (product: 250001). The assay used the QIAcuity 3x EvaGreen PCR Master Mix (product: 250113) with manufacturer recommended reaction set-up and cycling conditions with an annealing temperature of 58℃ (Qiagen: HB-2791 01/2021). The plate was imaged at 200 ms with a gain of 1. All reactions had acceptable levels of partition occupancy and mean 95% confidence intervals ±1.9% . Results from the dPCR were back-transformed to copies per ng of input RNA, and adjusted values were log10-transformed for statistical comparisons. Results were analyzed for normality, homoscedasticity and tested using a non-parametric Wilcoxon rank sum test with continuity correction. To take into account possible treatment effects on *Buchnera* numbers, we also quantified *syeA* transcripts relative to transcripts of a control *Buchnera* gene *rne*, encoding ribonuclease E. Methods were the same as for *syeE*, using primers 5AL81_F and 5AL81_R^18^. Values for #*syeA* transcripts/#*rne* transcripts were normally distributed with equivalent variance, so we used a Welch 2-sample t-test.

### Embryo phenotypes after *syeA* knockdown

To determine if *syeA* knockdown causes defects in colonization or development, 7-day-old (4th instar) aphids, which contain developing embryos, were injected with control PNAs or anti-*syeA* PNAs. To examine phenotypes of embryos that were at the stage of colonization at the time of exposure, we sampled 7 days later. Birth occurs 8-10 days after Stage 7 (when *Buchnera* colonizes)^19–21^, and our sampling time was chosen so that embryos exposed at Stage 7 would be near the end of prenatal development when sampled.

We performed fluorescence *in situ* hybridization (FISH) to visualize *Buchnera* within the embryos. Aphids were dissected in 70% ethanol and fixed in Carnoy’s Solution (60% ethanol, 30% chloroform, 10% acetic acid) for 1 h then dehydrated in successive washes of 100% ethanol. Samples were then bleached in 6% hydrogen peroxide in 80% ethanol at room temperature for 1 week, with the solution replaced every two days. Fixed bleached samples were washed in 70% ethanol before rehydration in PBST. Fluorescent in situ hybridization was performed as previously described^22^, in buffer (20 mM Tris-HCl, 0.9 M NaCl, 0.01% SDS, 30% formamide) with DAPI and the *Buchnera*-specific probe ApisP2A-Cy5 (5′-Cy5-CCTCTTTGGGTAGATCC-3). Samples consisting of 28 treatment and 25 control mothers were imaged under a Nikon Eclipse Epifluorescence microscope. Each was scored for presence/absence of clearly deformed embryos.

### Lysosome quantification after knockdown

To examine the effect of *syeA* knockdown on lysosome production in maternal bacteriocytes, 4–5-day-old (3rd instar) aphids were injected as described above. After 24 h, injected aphids were dissected in cold PBS, and bacteriocytes from individual aphids were placed into 0.2 mL tubes with 1X LysoView640 (Biotium) and DAPI. Samples were incubated for 30 min and imaged under a Nikon Eclipse Epifluorescence microscope. Fluorescence intensity was quantified per bacteriocyte.

### TEM to visualize flagellar basal bodies

The protocol followed that published previously^23^. Bacteriocytes of 7-day-old *A. pisum* str. LSR1were dissected into Buffer A, homogenized with a pestle, centrifuged at 4°C for 20 min, resuspended in Buffer A and kept on ice. A drop of the suspension was transferred to a carbon film-coated 400 mesh copper grid and allowed to be absorbed for 1–3 min, and dried with filter paper. Negative staining and imaging was performed immediately on a Jeol 1400Flash electron microscope.

---

**Extended Data Fig 1.**
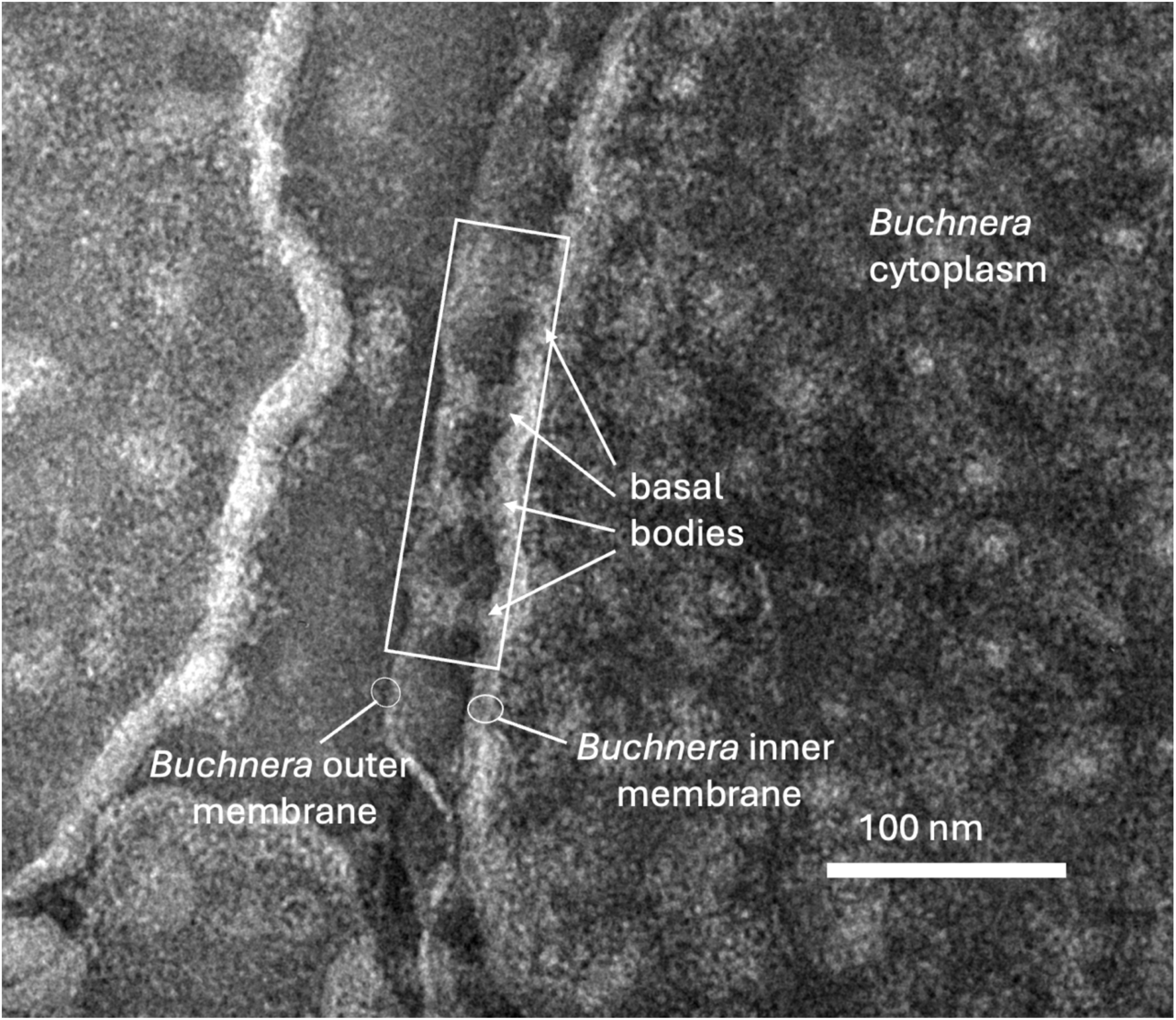
Negative stain TEM showing basal bodies of *Buchnera*-Ap. Scale bar 100 nm.

**Extended Data Fig 2.**
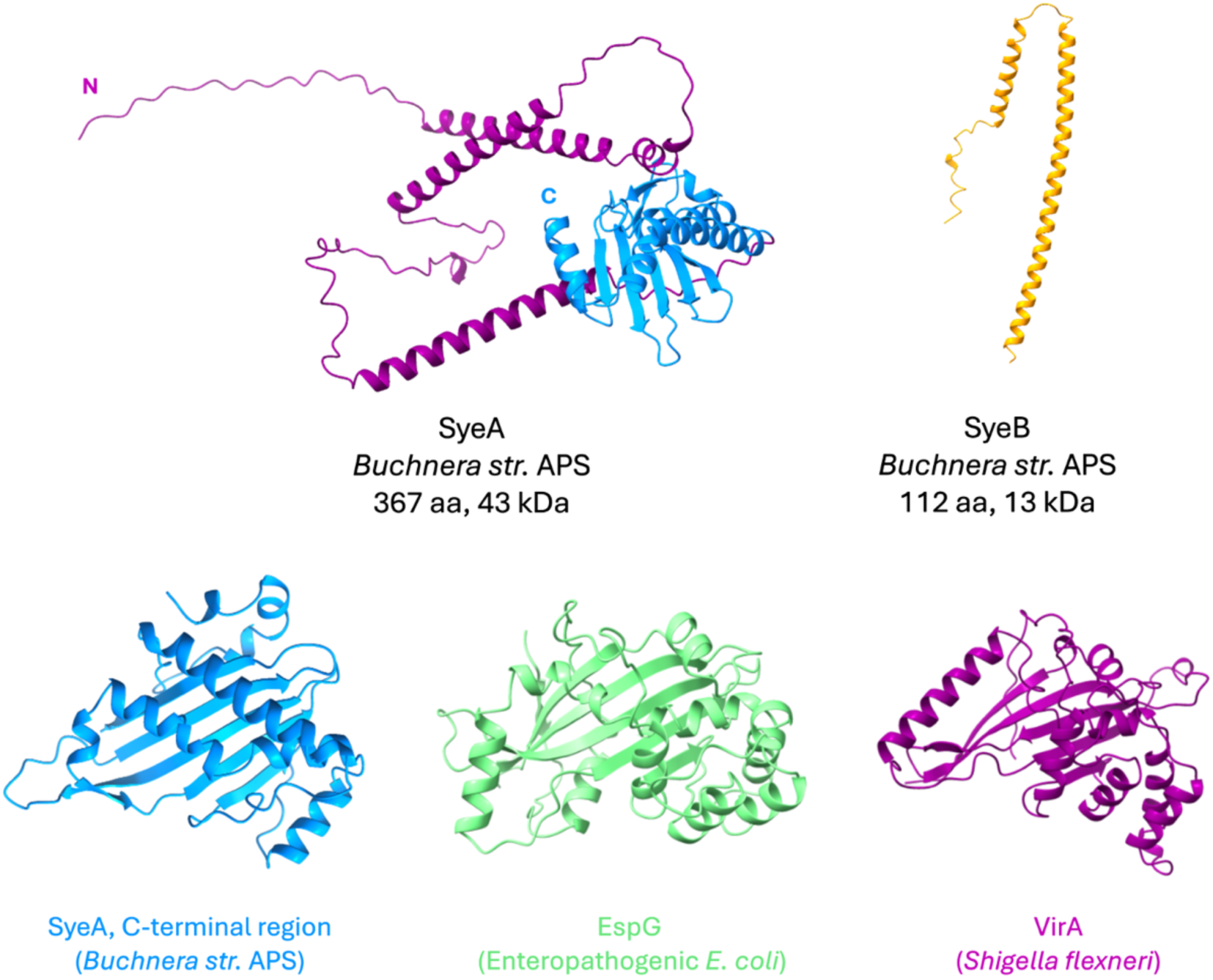
Structure of SyeA, SyeB, and homologs in pathogenic bacteria. Top: Full length *Buchnera*-Ap SyeA and SyeB. Bottom, left to right: conserved C-terminal region of SyeA of *Buchnera*-Ap, EspG of enteropathogenic *E. coli*, VirA of *Shigella flexneri.* Structures were predicted with AlphaFold3 and visualized with ChimeraX

**Extended Data Fig 3.**
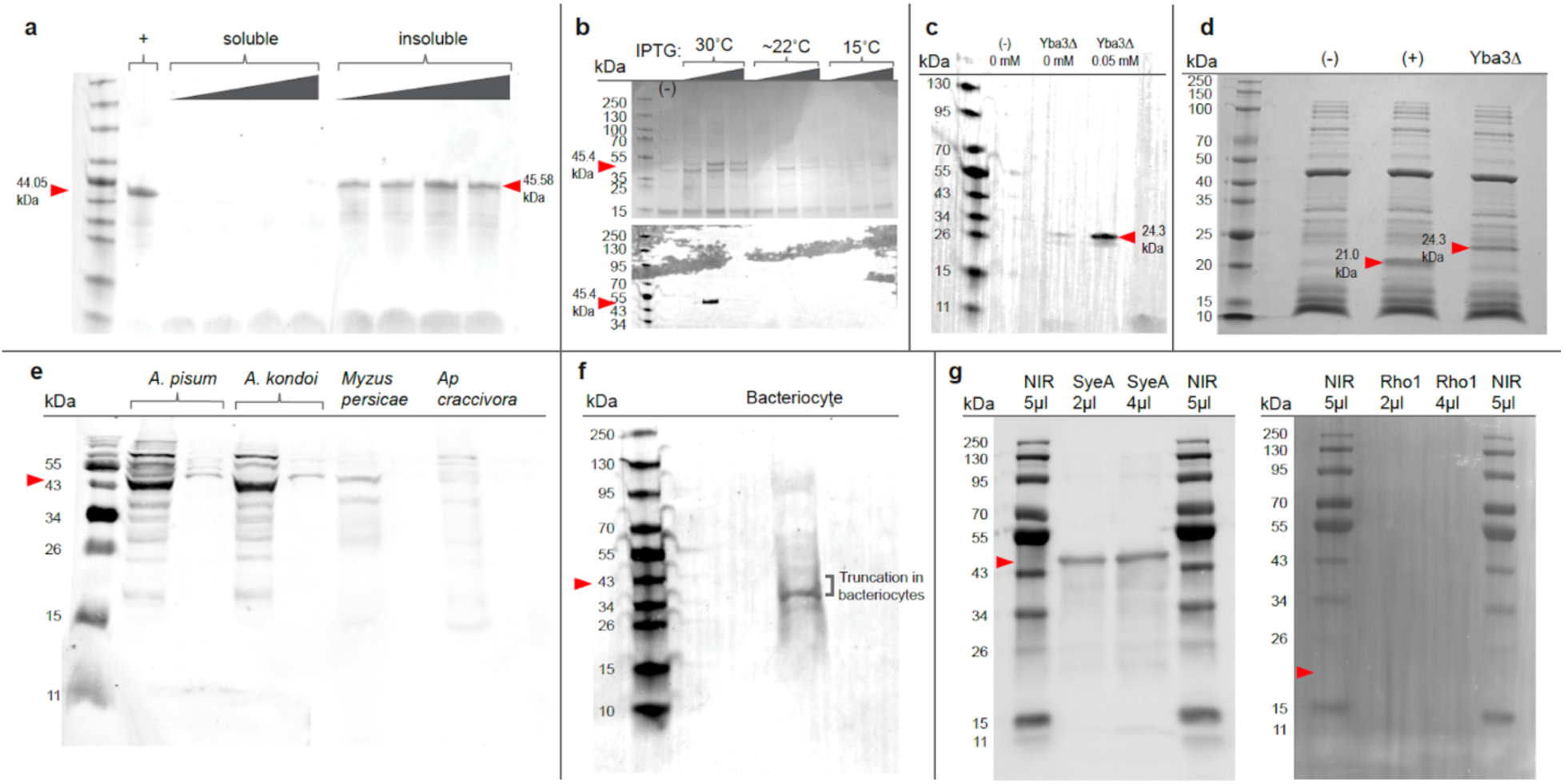
Detection of heterologously expressed and native SyeA. **a)** Western blot with anti-SyeA showing specificity and insolubility of SyeA expressed heterologously in *E. coli*. Positive control (+) was full-length SyeA (expressed and used for antibody production by Biomatik). The difference in molecular mass reflects the presence of a cleavage recognition site for removal of His6-tag. Soluble fractions are supernatants of pelleted culture following addition of lysis buffer and lysozyme. Insoluble fractions are from pellets remaining after lysis, denatured with urea and SDS. IPTG for induction of expression had concentrations of 0.0, 0.1, 0.5, and 1.0 mM. **b)** Upper: Coomassie staining of inclusion bodies following expression of full-length His6-SyeA at 30℃, ∼22℃, and 15℃. Lower: Western blot of **b**, using anti-His6 antibody. Negative control (-) was *E. coli* carrying the empty vector. Expected size of SyeA with the tag is 45.4 kDa. **c)** Western blot of inclusion bodies in *E. coli* carrying either the empty vector (-) or the His6-SyeAΔ1-172 vector, uninduced and induced at 0.05 mM IPTG. Expected size of SyeAΔ is 24.3 kDa. **d)** Expression of SyeAΔ with the PURExpress cell-free expression system. Lanes include reactions with no plasmid (-), control plasmid encoding *E. coli* dihydrofolate reductase (+), and plasmid expressing SyeAΔ. Expected size of (+) is 21 kDa. Expected size of SyeAΔ is 24.3 kDa. **e)** Western blot of lysate of whole bodies of 4 aphid species using anti-SyeA raised against a conserved peptide of SyeA, showing binding and specificity in *A. pisum* and *A. kondoi*, but not in two more distantly related species. Expected size in both is 43.3 kDa. **f)** Western blot of mature bacteriocyte samples following direct solubilization and boiling in 6X SDS dye, using anti-SyeA as the primary antibody at a concentration of 1:10,000. Expected size of SyeA in *Buchnera-*Ap is 43.3 kDa, indicating truncation to ∼34 kDa. **g)** Westerns showing lack of cross-binding by secondary antibody for Rho1 with anti-SyeA primary antibody. and Rho1. All 8 lanes were from the same gel, probed with anti-SyeA, then transferred to a membrane which was cut in half, with one half (left) probed with the anti-rabbit secondary antibody for anti-SyeA, the other half (right) with the anti-mouse IgG1 secondary for anti-Rho1.

**Extended Data Fig 4.**
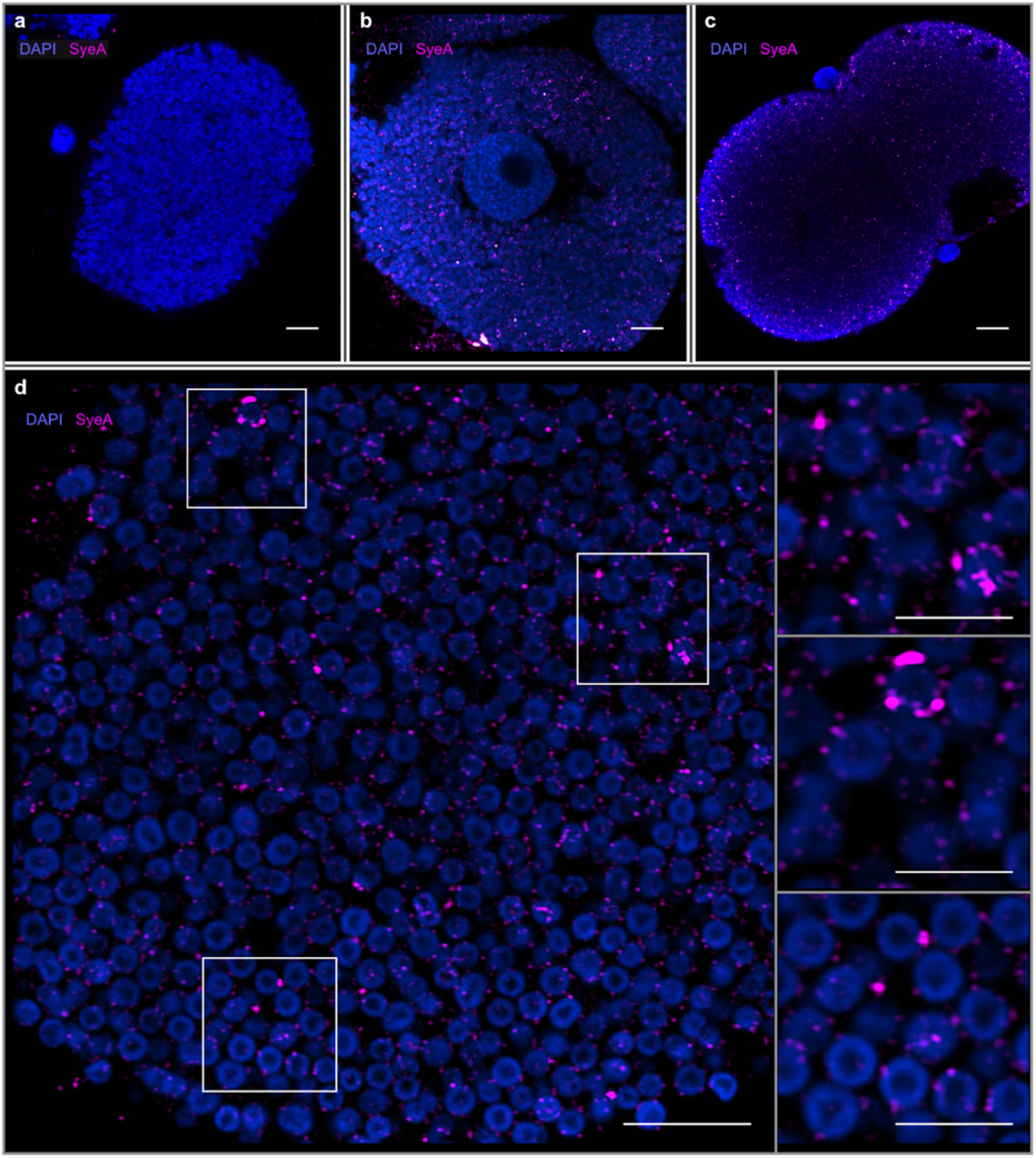
SyeA localization and abundance in mature bacteriocytes. (**a**) Representative negative control (no primary antibody and identical settings to **b, c**). (**b, c**) bacteriocytes from 7 d and 22 d old aphids. **d**) *Buchnera* cells within mature bacteriocyte showing location of SyeA outside *Buchnera* cells or diffused within symbiosomal compartments. Scale bars are 10 μm in **a**-**d**, 5 μm in insets of **d**.

**Extended Data Fig 5.**
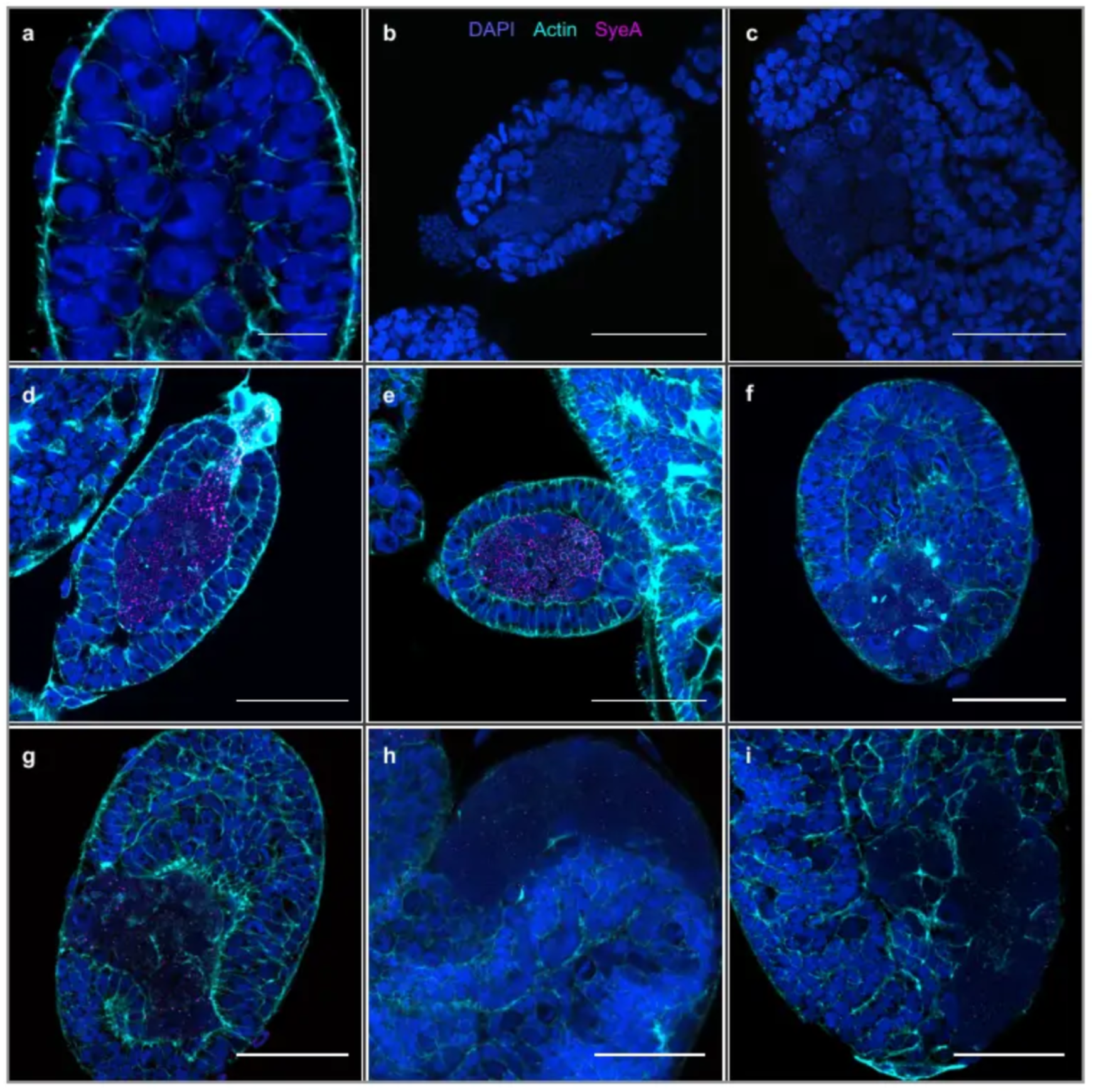
SyeA in embryos before and after division of syncytium into individual bacteriocytes. **a**) Control Stage 4 embryo prior to *Buchnera* colonization and lacking SyeA signal (fully stained, same settings as **b-i**)**. b-c)** Stage 8 and Stage 12 embryos with DAPI and no primary antibodies (no actin stain in b-c). **d**) Stage 7 embryo showing SyeA throughout multinucleate syncytium. **e**) Stage 8 embryo with SyeA forming rings at the symbiosome borders. **f**-**i**) Stages 11, 12, 13, and 14 embryos showing reduced and localized SyeA after formation of individual bacteriocytes. Scale bars: 10 μm in **a**-**c**, 50 μm in **d**-**i**.

**Extended Data Fig 6.**
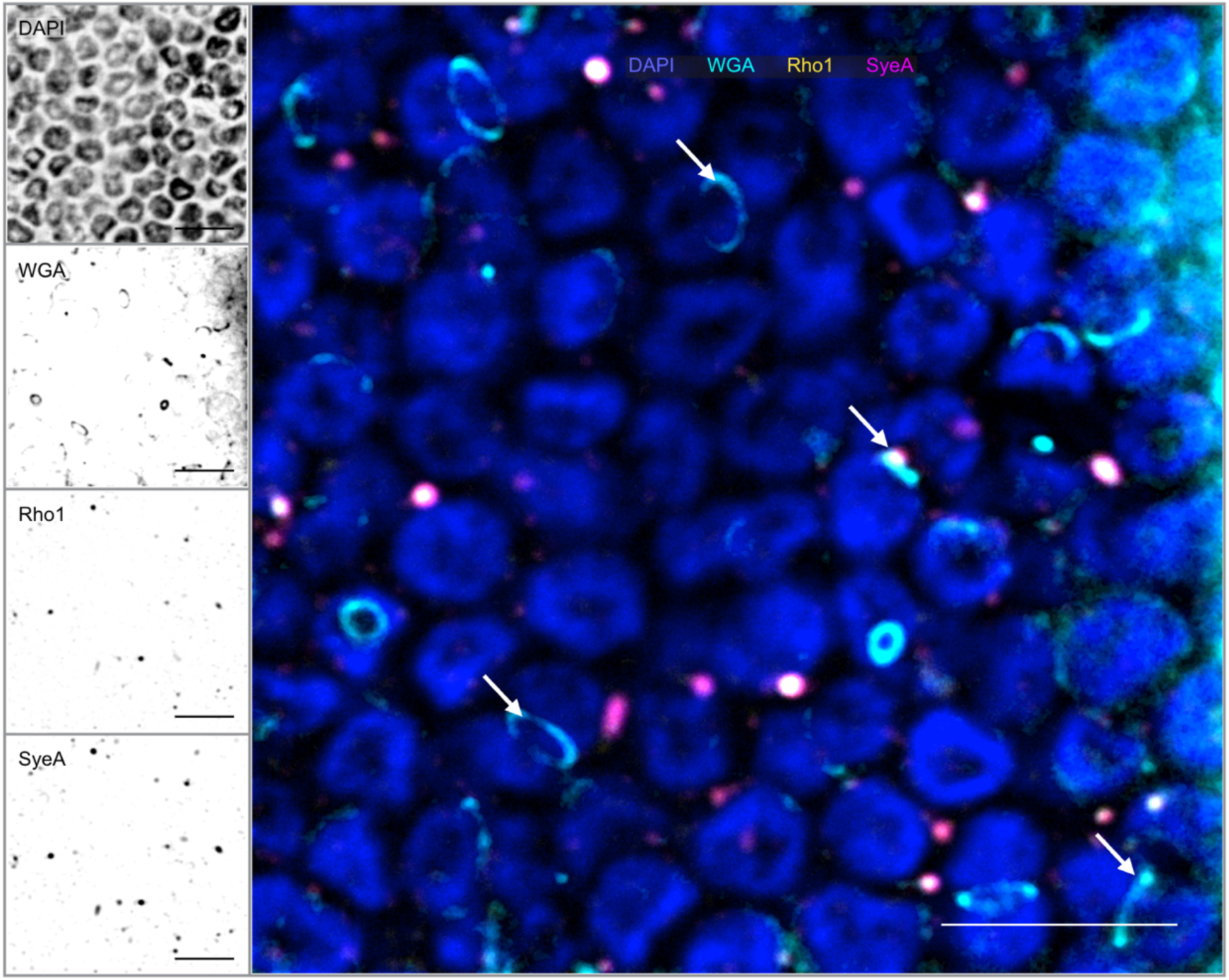
Colocalization of SyeA and Rho1 in mature bacteriocyte of 8-d-old pea aphid. WGA signal is strongest for the peptidoglycan layer forming at the septum of dividing *Buchnera* cells (examples indicated by arrows) and relatively weak for the cell wall perimeter. Scale bars 5 μm.

**Extended Data Fig 7.**
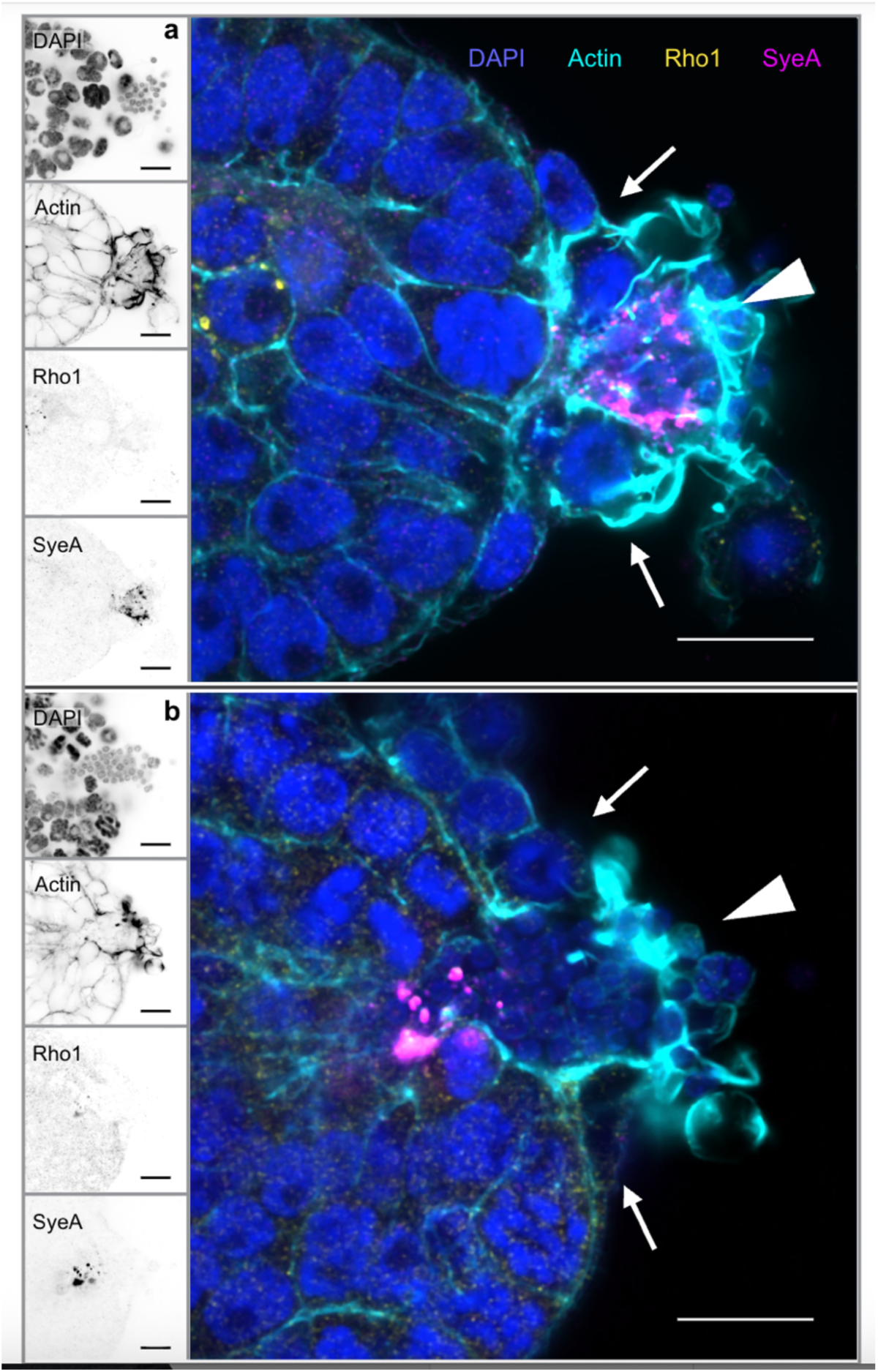
Actin polymerization at *Buchnera* entry point during colonization. White arrowheads indicate the entry point, which forms a channel through the center of the embryo. Phalloidin staining shows large actin mass around the extruded region of the syncytium, which extrudes beyond the cuticle of the embryo and is flanked by large follicle (maternal) cells (arrows). Scale bars 10 μm.

**Supplementary Table 1.**
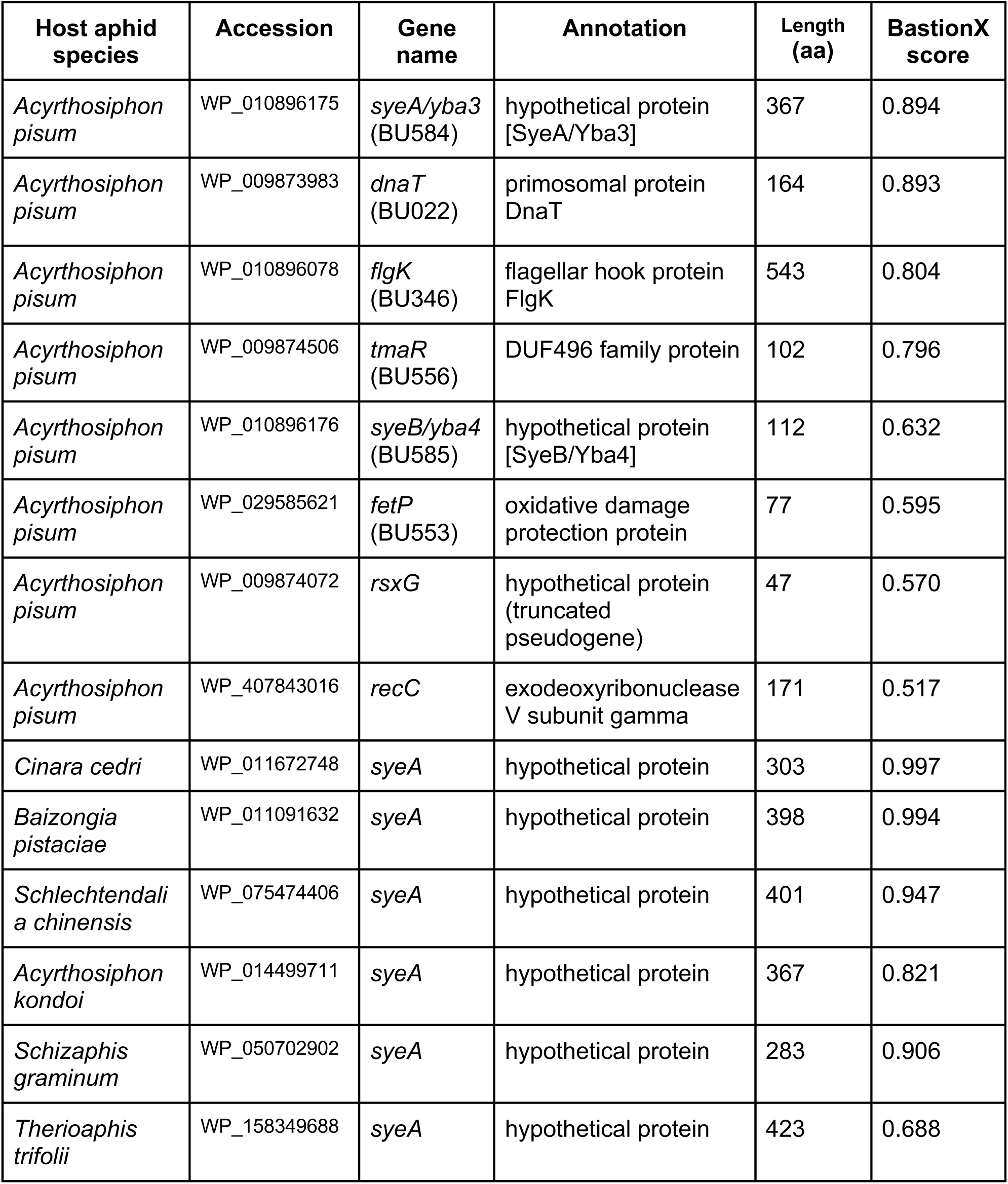
Predicted T3SS substrates from a screen of the complete proteome of *Buchnera*-Ap* and of SyeA homologs of *Buchnera* of representative aphid species of Fig 1b.

